# *Acanthamoeba castellanii* as a model for unveiling *Campylobacter jejuni* host-pathogen dynamics

**DOI:** 10.1101/2024.05.24.595673

**Authors:** Fauzy Nasher, Burhan Lehri, Richard Stabler, Brendan W. Wren

## Abstract

The persistence of the major enteric pathogen *Campylobacter jejuni* in the natural environment, despite being microaerophilic, remains unsolved. Its survival in the natural atmospheric environment likely stems from several factors, including interactions with amoebae. *C. jejuni* transiently interacts with Acanthamoebae and this is thought to provide protection against unfavourable atmospheric conditions and subsequently prime the bacteria for interactions with warm-blooded hosts. Acanthamoebae play vital roles in microbial ecosystems by preying on bacterial species, some of which are clinically important. We analysed the whole transcriptome of *A. castellanii* infected with *C. jejuni* 11168H. Our findings provide evidence that infection of *A. castellanii* with *C. jejuni* triggers distinct and reproducible cellular responses. Upregulated genes were associated with protein synthesis, DNA damage and repair, gluconeogenic pathways, and protein folding and targeting, while downregulated genes were involved in calcium ion transport, osmotic stress response, energy reserve metabolic processes, and protein hydroxylation. From this data we characterized Cj0979c, named here *C. jejuni* endonuclease (CjeN), which induces DNA damage in *A. castellanii*. High-resolution microscopy revealed an unexpected association between *C. jejuni* and host mitochondria, while infected cells show elevated cytosolic calcium levels and metabolic changes favouring “Warburg-like” metabolism. The increased lactate production was subsequently depleted, suggesting that this host metabolic by-product may support *C. jejuni* survival. These findings identify an unexpected interaction between amoebae and a microaerophilic bacterium and provides a useful model for further research on host-pathogen interactions.

## Introduction

The long-term relationship between amoebae and bacteria holds significant implications for the evolution of pathogens and has been termed the eco-evo perspective of bacterial pathogenesis [1, 2]. Amoebae are abundant unicellular organisms found in diverse environments that predate bacteria, influencing the evolution of both host and prey[3]. Free-living amoebae, such as Acanthamoebae, play essential roles in aquatic and terrestrial ecosystems, shaping microbial communities and nutrient cycling dynamics [4]. Notably, Acanthamoebae exhibit two distinct life cycles: an active phagotrophic trophozoite stage and a low-metabolic, resilient cyst stage, aiding survival under adverse conditions [5].

The co-evolution of bacteria and amoebae often precede bacterial interactions with warm-blooded hosts [6], with amoebae evolving mechanisms to ingest and kill bacteria, while bacteria develop strategies to resist predation and, at times, kill amoebae hosts [4, 6]. This microbial dynamic mirrors the intricate interplay observed between phagocytes of the warm-blooded host’s immune system and pathogenic bacteria [7].

*Campylobacter jejuni*, a microaerophile bacterium, challenges conventional understanding by persisting in the environment outside avian and mammalian hosts [8]. *C. jejuni* is the most frequent cause of bacterial gastrointestinal infection globally [9]. Despite being a leading cause of bacterial gastroenteritis, *C. jejuni* is rarely transmitted between humans and its high prevalence in human infection is difficult to explain. While the consumption and handling of poultry contribute significantly to human infections [10], the transmission routes to poultry from the environment remain unclear. This gap in our understanding of the prevalence and transmission of this major pathogen has been referred to as the “Campylobacter conundrum” [11]. Amoebae like Acanthamoebae have been proposed as a natural reservoir for Campylobacter, providing a niche and protection against potential toxic atmospheric conditions [12–14]. Acanthamoebae transiently host *C. jejuni*, priming the bacterium and facilitating its subsequent invasion of human epithelial cells and re-invasion of amoebae [15, 16]. Therefore, exploring Acanthamoebae-*C. jejuni* interactions offers a promising model for studying the adaptation, persistence, and infection dynamics of this enigmatic pathogen.

In the present study, we investigate the intricate interplay between Acanthamoebae and *C. jejuni*, aiming to elucidate the dynamics of their interactions and gain new insights into microbial pathogenesis mechanisms. Our findings reveal that the interplay between *A. castellanii* and *C. jejuni* triggers a series of cellular responses, as supported by experimental data and comprehensive genome-wide transcriptome analysis. Gene ontology (GO) term enrichment analyses highlights significant alterations in gene expression, affecting various biological, molecular, and cellular processes. Specifically, upregulated genes are associated with crucial functions such as protein synthesis, DNA replication and repair, metabolic activities, and protein folding and targeting. Conversely, downregulated genes involve processes including calcium ion transport, osmotic stress response, cytoskeleton organization, energy reserve metabolic processes, and protein hydroxylation. Notably, Cj0979c (now named <u>C</u>ampylobacter jejuni <u>e</u>ndo<u>n</u>uclease, CjeN), a putative secreted endonuclease induced DNA damage in *A. castellanii*. High-resolution microscopy revealed an unexpected association of *C. jejuni* with host mitochondria, concurrently, infected *A. castellanii* exhibit increased cytosolic calcium levels. Infected amoebae cells also displayed increased aerobic glycolysis and gluconeogenesis, indicative of metabolic rewiring and shift towards Warburg-like metabolism for immediate energy production. Additionally, this metabolic rewiring increased lactate production, recently shown to support *C. jejuni* population expansion during acute infection [17]. This host metabolic by-product potentially supports *C. jejuni* survival.

These results provide valuable insights into the molecular mechanisms governing the interaction between Acanthamoebae and *C. jejuni*, contributing to our understanding of microbial evolution and pathogenesis. The study represents a significant advancement in our understanding of the interaction between *C. jejuni* and Acanthamoebae and provides compelling evidence that *C. jejuni* manipulates the amoeba host to facilitate its survival, marking this a pivotal discovery in the field.

## Results

### *A. castellanii* differentially regulates its genes in response to *C. jejuni* infection

To uncover the response of *A. castellanii* to intracellular *C. jejuni*, we conducted whole-genome transcriptome analyses of *A. castellanii* at 4 hrs post-infection with *C. jejuni* 11168H. There was a significant difference (*p*<0.05) in gene expression between *A. castellanii* infected with *C. jejuni* and the control; a total of 196 genes were differentially transcribed (>2-fold; *p*-value < 0.05): 26 were up-regulated whilst 170 were down-regulated **(Supplementary File 1)**.

To verify our RNA-Seq results, RNA from *A. castellanii* harbouring *C. jejuni* was extracted independently of the RNA-Seq experiments for real-time RT-PCR. Genes that were verified are highlighted in **Figure 1a**. Upregulated genes encode heat shock protein 20 (*hsp20*) (XP_004336746.1), an ATP-independent chaperone that was shown to be associated with encystation [18]; a copper transport accessory protein (XP_004349665.1), possibly involved in copper homeostasis; a kinetochore like protein (XP_004334023.1), Kinetochore-like proteins are not well-defined, however, kinetochores themselves are protein complexes that assemble on chromosome during cell division, DNA repair and cell signalling [19] and finally we verified upregulation of a gene encoding for universal stress domain containing protein (USP) (XP_004336998.1), USP may provide a general “stress endurance” activity and enhance survival during prolonged exposure to stress [20].

**Figure 1:**
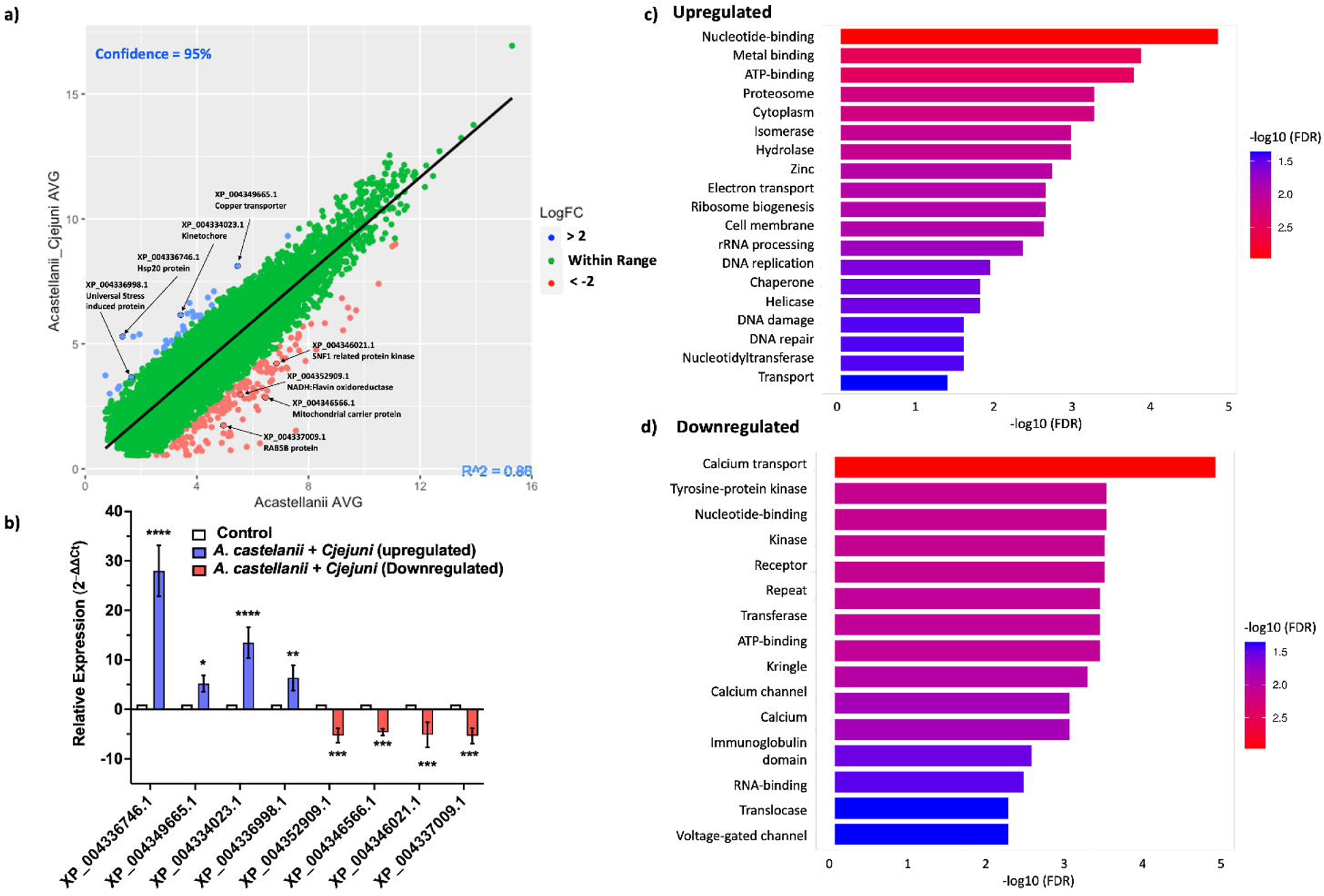
Validation and Enrichment Analysis of RNA-Seq Data. **a)** A scatter plot displaying the average fold change between *A. castellanii* infected with *C. jejuni* versus *A. castellanii* alone 4 hr post infection. Upregulated genes are indicated in blue, while downregulated genes are shown in red. **b)** Real-time RT-qPCR analysis of *A. castellanii* infected with *C. jejuni* 11168H. Upregulated genes encoding XP_004336746.1 (heat shock protein 20 (hsp20)), XP_004349665 (copper transport accessory protein), XP_004334023.1 (kinetochore-like protein), and XP_004336998.1 (universal stress domain-containing protein), are depicted in blue bars. Downregulated genes encoding XP_004352909.1 (NADH:flavin oxidoreductase), XP_004346566.1 (mitochondrial carrier protein), XP_004346021.1 (Sucrose non-fermenting 1-related protein), and XP_004337009 (RAB5B protein), are highlighted in red. Significance levels are indicated as follows: \**p*<0.05, \*\**p*<0.001, \*\*\**p*<0.0001. Error bars represent standard deviation (SD). **c)** Bar plot representing the enrichment of upregulated Gene Ontology (GO) terms. **d)** Bar plot illustrating the enrichment of downregulated GO terms. Enriched terms are indicated with colours corresponding to the significance level (-log10(FDR)). Plots were generated using ShinyGO 0.80 [24].

For downregulated genes, we verified genes encoding NADH:flavin oxidoreductase (XP_004352909.1), a free flavin reductase by NADH; mitochondrial carrier protein (MCP) (XP_004346566.1), MCPs transport nucleotides, amino acids, carboxylic acids, inorganic ions, and cofactors across the mitochondrial inner membrane, thereby connecting metabolic pathways of the cytoplasm with those of the mitochondrial matrix [21]. Sucrose non-fermenting 1-related protein (SNF1) (XP_004346021.1), SNF1 kinases are widely attributed to controlling various nutrient-responsive and cellular developmental processes within the cell [22]; and lastly RAB5B protein (XP_004337009.1), RAB5B is a Ras-related protein that is involved in early endosome movement, and is necessary for the fusion of vesicles [23].

Real-time RT-qPCR analysis revealed a statistically significant (p < 0.05) upregulation in the expression levels of genes encoding XP_004336746.1 (∼28-fold); XP_004349665.1 (∼5 fold); XP_004334023.1 (∼12-fold); XP_004336998.1 (∼7-fold) and the downregulation of genes encoding XP_004352909.1 (∼5-fold); XP_004346566.1 (∼4-fold); XP_004346021.1 (∼5 fold); and finally, XP_004337009.1 (∼5-fold) **(Figure 1b)**.

To further understand our RNA-Seq data, we conducted an enrichment analysis of both upregulated **(Figure 1c)** and downregulated **(Figure 1d)** genes. Gene Ontology (GO) enrichment analyses revealed several significantly enriched GO terms associated with biological processes, molecular functions, and cellular components. Of the upregulated biological processes, nucleotide-binding pathways were enriched, these included genes associated with DNA replication, nucleotide excision repair, and RNA processing. Enrichment of metal-binding activities suggested ion homeostasis, potentially regulating metal ion concentrations within the cell. We also observed enrichment of hydrolase, indicating processes such as protein degradation, lipid metabolism, and cellular signalling were upregulated. Enrichment in DNA damage and repair processes suggested that the upregulated genes may play crucial roles in maintaining genomic integrity and responding to DNA damage-induced stress during *A. castellanii* interactions with *C. jejuni*. Among the molecular functions, enrichment in ATP-binding activities suggested roles in energy metabolism, cellular transport processes, and ATP-dependent molecular pathways. Enrichment of the proteasome indicated protein degradation and regulation of cellular processes **(Supplementary File 3 (Figure S1))**. Upregulated cellular components enrichment included cytoplasmic components, this suggests increased cellular metabolic activity and protein synthesis within the cytoplasmic compartment. GO terms associated with cell membrane were also enriched, which indicated upregulation of gene involved in membrane structure, transport, and signalling processes **(Figure 1c)**.

Downregulated biological function genes included calcium transport, an indication of calcium homeostasis disruptions. Decreased Tyrosine-Protein Kinase activity implied impaired cellular signalling pathways and potential disruptions in signal transduction cascades and phosphorylation-mediated regulatory mechanisms. Downregulation of Nucleotide Binding suggested potential alterations in nucleotide metabolism, cellular signalling, and ion transport processes. Pathways involved in molecular functions included receptor activity, which suggested potential impairments in cellular communication and responsiveness to external stimuli. Transferase Activity were also downregulated, an indication that potential reduction in protein modification, lipid metabolism, and signal transduction. We also observed downregulation of ATP Binding, this suggests alterations in energy-dependent cellular processes. Downregulation of Cellular Component pathways included genes associated with Repeat Regions, this may indicate alterations in protein domains or structural motifs crucial for protein-protein interactions and functional diversity. Calcium channel genes were also enriched, suggesting disrupted calcium ion influx, which is essential for intracellular signalling. Immunoglobulin domain genes were also enriched. Reduced expression of genes containing immunoglobulin domains suggested potential impairments in immune responses and antigen recognition processes. Translocase Activity was also reduced, indicating alterations in membrane transport (import/export) processes and organelle biogenesis. Voltage-Gated Channel was also downregulated, indicating disruptions in ion transport. Finally, downregulation of RNA binding genes suggests potential impairments in RNA processing, stability, and regulation of gene expression **(Figure 1d)**. All enriched pathways and fold enrichment are provided in the supplementary material **(Supplementary file 2)**.

### CjeN *(cj0979c),* a *C. jejuni* endonuclease induces double-stranded breakages (DSBs) during infection

Our RNA-Seq data indicated that *A. castellanii* infected with *C. jejuni* upregulated pathways that are involved in DNA damage and repair. To confirm this, we used anti-γHA2.X as a biomarker for DNA damage and monitor DNA double stranded breakages (DSBs) (**Figure 2**).

**Figure 2:**
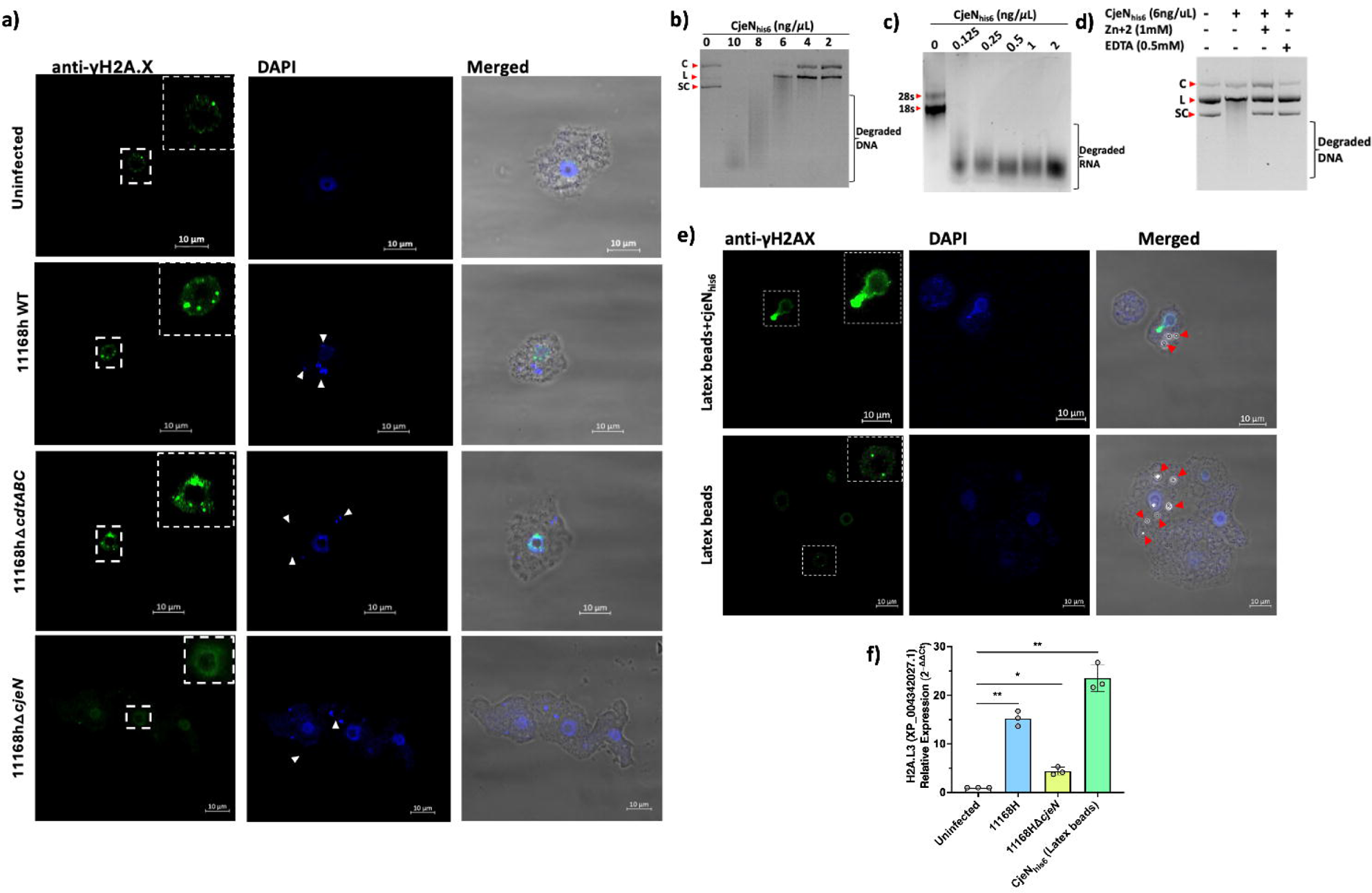
Mechanisms of DNA damage in *A. castellanii*. **a)** Anti-γHA2.X biomarker of double stranded breakages (DSBs) revealed *C. jejuni* 11168H wildtype, 11168HΔ*cdtABC* and 11168HΔ*cjeN*+*cjeN* mutant cause multiple DNA damage at 4 hrs post infection relative to uninfected cells and 11168HΔ*cjeN.* **b)** CjeN nuclease activity on plasmid double stranded DNA, smear indicate degraded DNA (C; circular, L; linear, SC; supercoiled). **c)** CjeN activity of eukaryotic RNA (28s and 18s indicate the type of RNA). **d)** Effect of Zinc (Zn^2+^) or ethylenediaminetetraacetic acid (EDTA) on CjeN nuclease activity. **e)** Effect of mutation to the active site of CjeN (CjeN_Thr69Ser77Thr111his6)_ on nuclease activity **f)** CjeN adsorbed onto latex beads and its activity was monitored *in vivo*. **g)** Relative expression of H2A histone family member L3 (XP_004342027.1) of *A. castellanii* infected with *C. jejuni* 11168H wildtype, its Δ*cjeN* mutant or recombinant CjeN_hi6_ protein adsorbed onto latex beads by RT-qPCR at 4 hr post infection. Significance levels are indicated as follows: \**p*<0.05, \*\**p*<0.001. Error bars represent standard deviation (SD). For images: The amoebae are visible in the transmitted-light channel (labelled: Merged); Nucleic acid and *C. jejuni* cells are visible in blue channel (labelled: DAPI Ex/Em 350/465 nm); Anti-γHA2.X in the green channel (labelled: Anti-γHA2.X Ex/Em 493/518 nm). Arrows indicate bacteria (white) or latex beads (red); images are representation of three independent biological replicates; images were captured with ×63 oil objective Scale bar: 10Lμm.

Our findings revealed that *C. jejuni* 11168H causes multiple DSBs to *A. castellanii* DNA **(Figure 2a).** Similar to other Gram-negative pathogens, *C. jejuni* synthesizes a cytolethal distending toxin (CDT), which was reported to cause direct DNA damage [25], thereby activating DNA damage checkpoint pathways. Comprising three protein subunits, CDT includes CdtA, CdtB, and CdtC, with CdtB serving as a nuclease. To elucidate the mechanism by which *C. jejuni* induces DSBs in *A. castellanii*, we generated a defined *cdtABC* deficient mutant of *C. jejuni* 11168H **(Supplementary File 3 (Figure S3))**. Subsequently, we infected *A. castellanii* with this 11168HΔ*cdtABC* mutant and DSBs (γHA2.X) were monitored **(Figure 2a)**. Intriguingly, despite the absence of the *cdtABC* genes, multiple DSBs were still observed in *A. castellanii* infected with the *C. jejuni* 11168HΔ*cdtABC* mutant. This suggested that *C. jejuni* 11168H employed other mechanisms to induce DSBs.

Previously, our RNA-Seq analyses of intra-amoebic *C. jejuni* showed upregulation of an uncharacterized gene, *cj0979c*, which was predicted to be a secreted endonuclease [16]. Here, a protein BLAST search (https://blast.ncbi.nlm.nih.gov/) revealed that *cj0979c* shares identities with *Staphylococcus aureus* Nuc1 (32.80%) and *Bacillus subtilis* 168 YncB (31.30%) proteins **(supplementary File 3 (Figure S2))**. Both Nuc1 and YncB are sugar non-specific extracellular endonucleases [26]. To explore the role of *cj0979c* (now designated as *<u>C</u>ampylobacter jejuni* <u>e</u>ndo<u>n</u>uclease, CjeN) in inducing double-strand breakages, we generated *C. jejuni* 11168HΔ*cjeN* deficient strain and infected *A. castellanii* with this mutant. Interestingly, no DSBs (γHA2.X) were detected in *A. castellanii* infected with this mutant strain, this phenotype was restored when *cjeN* was complemented with a copy in the rRNA gene clusters on the chromosome **(Figure 2a)**.

We sought to characterize CjeN, we therefore expressed and purified recombinant CjeN_his6_ that lacked the first 39 amino acids (predicted signal peptide (SignalP 5.0)) [27]. Our results confirmed that CjeN is a nuclease and was capable of digesting double stranded DNA (plasmid DNA) and RNA non-specifically in the presence of 130 µM calcium (Ca^2+^) and 5 mM magnesium (Mg^2+^) **(Figure 2b and c)**. Notably, CjeN digestion of DNA produced linear form of the plasmid DNA, a characteristic of endonucleases. Furthermore, CjeN activity was found to be inhibited by Zinc (Zn^2+^) or EDTA, a metal chelator **(Figure 2d)**. Zn^2+^ is known to hinder Ca^2+^ and Mg^2+^ dependent endonuclease activity [28] by competing for the catalytic site. Multiple sequence analysis (MSA) of CjeN, Nuc1 and YncB identified conserved active sites (CjeN; Arginine69, Glutamic acid77 and Arginine111 **(Figure S2)**), we generated a recombinant protein with substitution to all three sites, CjeN_Thr69ser77Thr111his6,_ and confirmed that mutation to these sites affects CjeN nuclease activity **(Figure 2e)**. Next, we sought to confirm CjeN endonuclease activity *in vivo* in the absence of *C. jejuni*. We immobilized recombinant CjeN_his6_ onto latex beads and incubated approximately 0.3 ng of the protein with *A. castellanii*. Subsequently, we observed double-stranded breakages (DSBs) indicated by γHA2.X, suggesting that CjeN was likely responsible for inducing DNA damage in *A. castellanii* host **(Figure 2f)**. Additionally, through RT-qPCR analysis, we found that the relative expression of *A. castellanii* H2A histone family member L3 (XP_004342027.1) was upregulated in the presence of both *C. jejuni* 11168H and recombinant CjeN_his6_, consistent with our RNA-Seq analyses **(Figure 2g)**.

### *C. jejuni* associates and alters host mitochondria dynamics

We observed GO terms related to mitochondrial function were enriched, including GO:0005759 (mitochondrial matrix), GO:0009055 (electron carrier activity), GO:0022900 (electron transport chain), GO:0006099 (tricarboxylic acid cycle), GO:0005524 (ATP binding), and GO:0016887 (ATPase activity). Conversely, there was downregulation of GO:0000281 (mitochondrial membrane). Given this, we decided to further investigate mitochondria **(Figure 3)**.

**Figure 3:**
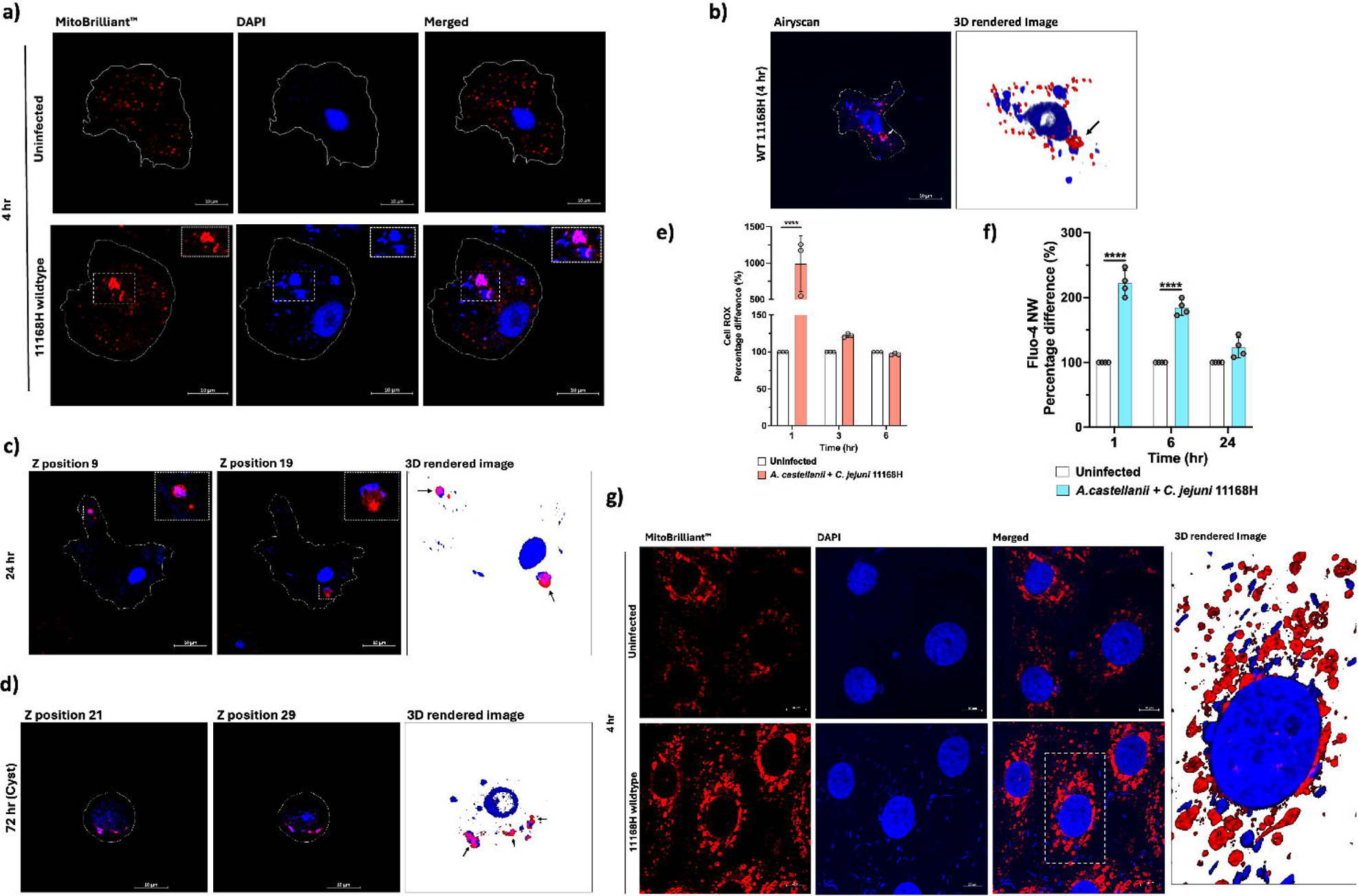
Mitochondrial Alterations, ROS, and Calcium Dynamics in *A. castellanii* Infected with *C. jejuni.* **a)** Confocal microscopy images comparing mitochondria in uninfected *A. castellanii* (above) with those infected with *C. jejuni* for 4 hrs (below). Mitochondria exhibit uniform distribution in uninfected cells, while infected cells show altered morphology with fused mitochondria. **b)** Volume view of confocal Z-stack image showing *A. castellanii* infected with *C. jejuni*, highlighting the altered mitochondrial morphology**. c)** Z-stack positions illustrating *A. castellanii* mitochondria dynamics at 24 hrs post *C. jejuni* infection, revealing fused mitochondria closely associated with *C. jejuni*-containing vacuoles. **d)** Z-stack positions depicting *A. castellanii* cysts at 72 hrs post *C. jejuni* infection, demonstrating persistence of altered mitochondrial morphology within cysts. **e)** Measurement of reactive oxygen species (ROS) using CellROX™ Deep Red, with pale pink bars representing *A. castellanii* infected with *C. jejuni* and white bars representing uninfected cells. **f)** Measurement of cytosolic calcium (Ca^2+^) levels using Fluo-4 NW Calcium Assay. ROS and Ca^2+^ data are presented as percentage. CellROX™ Deep Red fluorescent was measured at (Ex 644/Em 655 nm) and Fluo-4 NW (EX 490/Em 525 nm). **g)** Images comparing mitochondria in uninfected human intestinal epithelial T84 cells with those infected with *C. jejuni* at 4 hrs. Mitochondria were stained with Mitobrilliant™ Red, with *C. jejuni* and nucleic acid labelled with DAPI in the blue channel (DAPI) and mitochondria visible in the red channel. Scale bar: 10Lμm; *A. castellanii* were imaged at 100x, and human T84 cells were imaged at 63x. All images are representation of three biological replicates. Significance levels indicated as \*\*\*\**p*<0.0001. Error bars represent standard deviation (SD).

We employed Mitobrilliant™ 646 (excitation 655/Emission 668), a mitochondria-specific stain that relied on mitochondria membrane potential (ΔΨm) for uptake and confocal microscopy with Airyscan to facilitate our analysis. *Acanthamoeba spp*. are recognized for possessing an exceptionally high number of mitochondria [29], typically distributed uniformly within the cell. Remarkably, in *A. castellanii* cells infected with *C. jejuni*, we observed a distinct alteration in mitochondrial morphology, with mitochondria appearing fused **(Figure 3a)**. Notably, these fused mitochondria exhibited close association with vacuoles containing bacteria **(Figure 3a and b)**. This phenomenon persisted even at 24 hrs post-infection **(Figure 3c)**, and strikingly, even within *A. castellanii* cysts containing *C. jejuni* at 72 hrs post-infection **(Figure 3d)**, indicating the continuity of this phenomenon throughout the infection process. Interestingly, this phenomenon of mitochondria association was not observed with *C. jejuni* 11168HΔ*ciaB*, a mutant that was trafficked into digestive vacuoles and killed at a faster rate relative to the wildtype and a +*ciaB* complement strain (*C. jejuni* 11168HΔ*ciaB*+*ciaB*) **(Supplementary File 3 (Figure S4))**. This implied that *C. jejuni* confined within non-digestive vacuoles are more likely to interact and associate with host mitochondria.

Mitochondria is a significant source of reactive oxygen species (ROS) within cells [30]. Therefore, we measured ROS levels at different time points to investigate whether *A. castellanii* mitochondria were trafficked to the *C. jejuni* containing vacuole in response to the infection **(Figure 3e)**. We observed a significant (*p*<0.05) ∼12-fold increase in ROS levels at 1 hr post infection with *C. jejuni*, coinciding with the early stages of *C. jejuni* internalization. Interestingly, ROS was dramatically decreased at 3 hr, returning to levels like those in uninfected cells. This pattern persisted at the 6 hr, suggesting a dynamic regulation of ROS production during infection. We concluded that other mechanisms likely contribute to the close association between the *C. jejuni* containing vacuole and *A. castellanii* mitochondria.

Mitochondria also play a crucial role in regulating cellular calcium levels, a process tightly controlled due to calcium’s role as a catalyst for regulatory cascades impacting various cellular functions [31]. Given that we observed significant decrease in transcripts related to calcium homeostasis in our RNA-Seq data **(Supplementary file 1 and 2)** and our observation of the unusual mitochondria phenotype during *C. jejuni* 11168H infection, this prompted us to measure intracellular calcium levels in *A. castellanii* **(Figure 3f)**. Significantly (p<0.05) increased calcium levels were observed at 1 hr (∼2.21-fold) and 6 hrs (1.84-fold) post-infection, while at 24 hrs we observed a non-significant (∼1.22-fold) elevation in calcium levels.

To investigate whether there is a parallel in mitochondrial alteration phenomenon in warm blooded hosts, we infected human intestinal epithelial T84 cell with *C. jejuni* 11168H and tracked mitochondria after 4 hrs of infection **(Figure 3g)**. Interestingly, we observed altered host mitochondria dynamics and intimate association with *C. jejuni*, similar to our observation of *A. castellanii* infected with *C. jejuni*. This underscores the profound impact of this pathogen on cellular architecture and also likely their function.

### *C. jejuni* induced *A. castellanii* metabolic rewiring

Recent evidence suggests that eukaryotic cells undergoing bacterial infection exhibit distinct metabolic rewiring compared to their uninfected counterparts [32]. Consistent with this, we noted a significant (p<0.05) upregulation in the expression of pivotal gluconeogenic genes encoding phosphoenolpyruvate carboxykinases (PEPCK), specifically XP_004346619.1 (∼2.46-fold) and XP_004346618.1 (∼2.20-fold), as indicated by our RNA-seq analyses **(Supplementary file 1)**. When RT-qPCR was conducted, we observed a 2.33-fold increase for XP_004346619.1 and a 3.16-fold increase for XP_004346618.1 **(Figure 4a)**.

**Figure 4.**
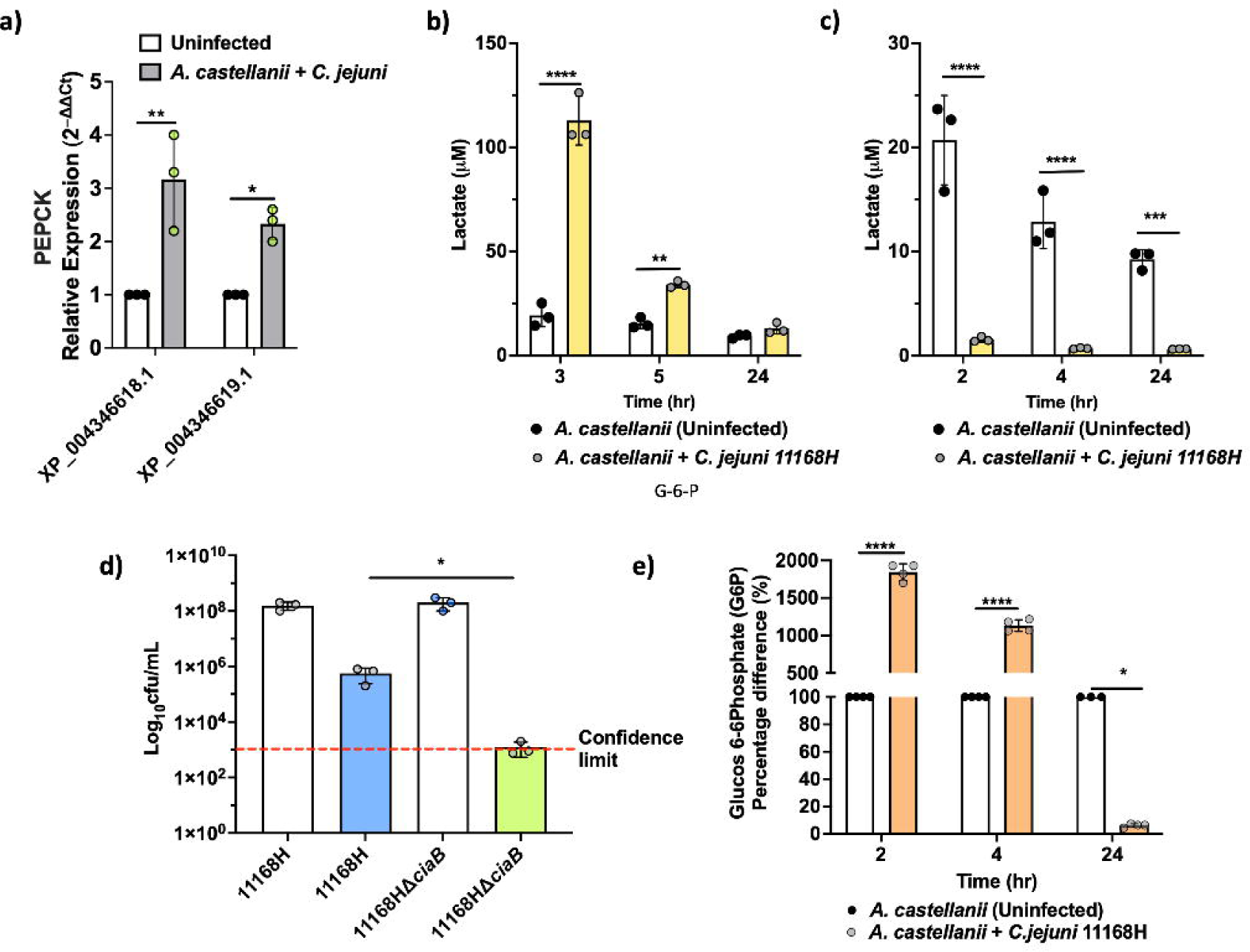
*A. castellanii* switches to Warburg-like metabolism at earlier time points of infection. **a)** Real time RT-qPCR of PEPCK encoding genes in *A. castellanii* infected with *C. jejuni* at 4 hr post infection. **b)** Extracellular lactate concentration 3, 5 and 24 hr post *C. jejuni* infection. *A. castellanii* were infected for 1 hour prior to 100 μg/mL gentamycin treatment. **c)** Extracellular lactate concentration 2, 4 and 24 hrs post *C. jejuni* infection (no gentamycin treatment). **d)** Intracellular survival of *C. jejuni* wildtype, Δ*lctP* mutant and Δ*lctP+lctP* after 4 hr infection (including 1 hr gentamycin treatment). **e)** Glucose 6 phosphate (G6P) concentrations in *A. castellanii* infected with *C. jejuni* compared to uninfected. Significance levels indicated as \**p*<0.05, \*\**p*<0.001, \*\*\**p*<0.0001, \*\*\*\**p*<0.0001. Survival is presented in Log scale. Error bars represent standard deviation (SD). SpectraMax Id5 was used to measure bioluminescence for lactate and G6P was measured by fluorescence (Ex540/Em 590 nm).

To investigate cellular metabolic activity, we monitored gluconeogenesis, the process of generating glucose from non-carbohydrate precursors, particularly during fasting, starvation, or low carbohydrate intake. As an indicator of glycolysis activity, we measured lactate production, a by-product of this metabolic pathway **(Figure 4b)**. We observed a notable increase in extracellular lactate in *A. castellanii* infected with *C. jejuni* 2 hrs post gentamycin treatment, with levels reaching ∼112 μM at 3 hrs post-infection, significantly higher than the ∼19 μM observed in uninfected cells. This heightened lactate production persisted after 5 hrs of infection, with infected *A. castellanii* maintaining elevated levels of lactate (∼34 μM), while uninfected cells remained relatively stable at ∼18 μM. Interestingly, lactate levels became comparable between infected and uninfected cells at the 24 hrs time point.

*C. jejuni* is capable of utilizing lactate from its environment [33], therefore, we measured lactate levels 2, 4 and 24 hrs post-infection before gentamicin treatment. We observed a significant reduction in lactate levels **(Figure 4c)** in *A. castellanii* with *C. jejuni* compared to uninfected *A. castellanii*. It is possible that under certain circumstances *A. castellanii* may enhance *C. jejuni* survival outside of the amoeba [34].

Previously, we reported a significant (p<0.05) upregulation of *lctP* (a putative lactate permease) in the *C. jejuni* 11168H transcriptome during intracellular survival within *A. castellanii* [16]. Here, mutation to *lctP* showed a significant (p<0.05) reduction in *C. jejuni* survival **(Figure 4d)**, although this did not lead to a complete failure in intra-amoeba survival suggesting other mechanism(s) of lactate transport into the cell [33, 35].

Additionally, glucose-6-phosphate (G6P) levels was assessed; G6P is the first intermediate of glucose metabolism and critical for glycolysis. G6P was substantially increased, reaching up to ∼18.45-fold at 2 hrs post-infection, and maintaining significant elevation (∼11.33-fold) at 4 hrs. Intriguingly, at 24 hours post-infection, we observed significantly decreased G6P levels (∼16-fold) in *A. castellanii* infected with *C. jejuni* 11168H compared to uninfected cells **(Figure 4e)**.

## Discussion

*Campylobacter jejuni* is the leading cause of bacterial foodborne gastroenteritis worldwide [36]. Investigating the dynamics between *C. jejuni* and Acanthamoebae is important as it illuminates the survival and persistence mechanisms of this pathogen in the environment outside its warm-blooded hosts. In this study, we investigated the response of *A. castellanii* to intracellular infection by *C. jejuni* using RNA-Seq.

The identified upregulated and downregulated transcripts shed light on the molecular mechanisms that underscore the interactions between *A. castellanii* and *C. jejuni*. Our findings revealed previously unreported interactions and highlight the complex interplay between host and pathogen at the molecular level. This study provides valuable insights into the strategies employed by *C. jejuni* to subvert host cellular processes. Upregulated processes such as DNA replication, cell cycle regulation, and protein folding suggested an active cellular response to infection, likely aimed at repairing DNA damage and maintaining cellular homeostasis. ATPase activity, electron transport chain components, and hydrolase activity indicated an increase in energy production and metabolic activity, possibly to support host defence mechanisms and cellular processes crucial for combating infection. On the other hand, the downregulation of processes related to calcium ion transport, peptidyl-tyrosine phosphorylation, and osmotic stress response may reflect the hijacking of host cellular machinery by *C. jejuni* to create a favourable intracellular environment for its survival. Based on this new data we provide a hypothetical model of the *C. jejuni* life cycle and its interaction with free-living amoeba **(see Figure 5)**.

**Figure 5.**
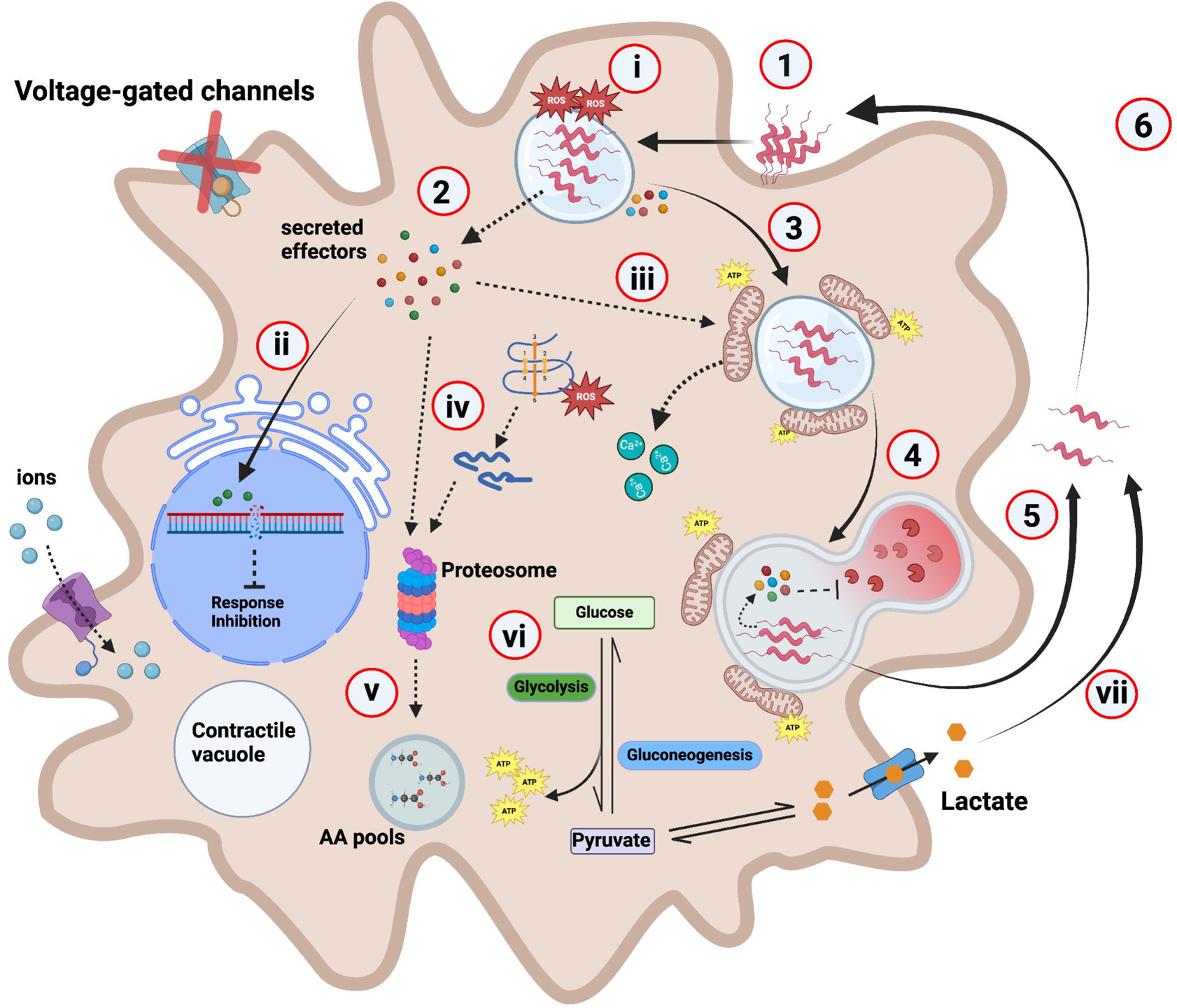
Hypothetical model depicting *C. jejuni* interaction with *A. castellanii*. 1. *C. jejuni* interacts with and enters *A. castellanii* through the aid of glycosylated FlaA [37] using an unknown receptor. **2.** Upon internalization, *C. jejuni* likely secretes effectors including campylobacter invasion antigens (Cias). **3.** Cias possibly promote *C. jejuni* trafficking and the formation of *C. jejuni* vacuoles. **4.** Indeed, a Δ*ciaB* mutant is digested and cleared at a faster rate relative to the wildtype **(Supplementary File 3 (Figure S3)). 5.** *C. jejuni* are capable of withstanding *A. castellanii* digestion and are exocytosed back into environment [16], **6.** where these “primed” *C. jejuni* can re-enter the Acanthamoebae transient host at a higher rate [15]. **i)** *A. castellanii* produces high levels of ROS to combat *C. jejuni* infection, this ROS level reduces to similar levels to the uninfected after 3 hr infection. **ii)** Effectors such as CjeN cause DNA damage, possibly damping immune response and inhibit other cellular responses. **iii)** We hypothesise that a yet to be identified effector possibly induces mitochondrial fusion in *A. castellanii*, leading to its relocation towards the periphery of the vacuole containing *C. jejuni*, possibly to provide macronutrients to the bacteria. Alterations to the mitochondria likely causes calcium influx into the cytosol. **iv)** *A. castellanii* upregulates proteasomal complex to degrade *C. jejuni* effectors as well as damaged endogenous proteins, **v)** this generates amino acids (AA) that feed into AA pools. **vi)** *C. jejuni* infection rewires *A. castellanii* metabolism, glycolysis and gluconeogenesis are increased, consequently, this increases ATP production. **vii)** Elevated glycolysis results in increased lactate as a by-product, which is then secreted where it is used by extracellular *C. jejuni*. Dashed lines indicate experimentally unproven pathways; cross on the voltage-gated channel indicates halted/or reduced activity.

*C. jejuni* endonuclease, CjeN, has been characterized in this study. Upon *C. jejuni* 11168H infection, CjeN induces double-strand breaks (DSBs) in *A. castellanii*, triggering DNA damage and repair response mechanisms. DSBs are particularly harmful lesions to DNA, requiring repair mechanisms to maintain chromosomal integrity [38]. Errors in this process can lead to mutations, and rearrangements of chromosomes, ultimately contributing to cell death [39]. Here, we present evidence that supports CjeN induces DSBs in *A. castellanii* during infection. CjeN degrades both double stranded DNA and RNA, and its activity was dependent on divalent metal ions Ca^2+^ and Mg^2+^, and Zn^2+^ inhibits its activity. Metal homeostasis is central to host-pathogen interactions [40]; therefore, it was not surprising to have observed differential regulation of genes associated with several metal ions, including those associated with Ca^2+^, Zn^2+^ and copper (Cu^2+^) homeostasis during *C. jejuni* infection of *A. castellanii*. Although CjeN has a signalling peptide and is likely transported to the extracellular environment, it remains unclear whether it is consistently cleaved and detached from the bacteria or if it remains attached to the surface in some instances, similar in other bacteria [41]. We propose that DSBs caused by CjeN may impair the ability of host cells to mount an effective immune response and interfere with cellular signalling pathways involved in responding to bacterial infections, allowing *C. jejuni* to persist for longer periods. CjeN was previously proposed to be involved in biofilm dispersal [42], it is quite likely that CjeN may also promote *C. jejuni* immune evasion during infection of warm-blooded hosts by degrading neutrophil extracellular traps (NETs) [43].

*C. jejuni* 11168H infection of *A. castellanii* induced alteration to host mitochondria, causing fusion and their relocation to the periphery of *C. jejuni* containing vacuoles. Mitochondria play a pivotal role in orchestrating various cellular processes and host defence mechanisms during bacterial infections. Numerous pathogens, *Legionella pneumophila* [44], *Mycobacterium tuberculosis* [45] and *Chlamydia spp*. [46] manipulate mitochondrial dynamics and functions to enhance their intracellular survival or evade immune detection. Mitochondria dynamic behaviour are characterized by continuous fission, fusion, and trafficking. These processes are dictated by the cellular energy demands, wherein mitochondrial fusion is favoured in conditions requiring increased ATP production via oxidative phosphorylation (OXPHOS) [47]. Consistent with this, we also observed an increased electron transport chain (ETC) and its activity in *A. castellanii* infected with *C. jejuni*, suggesting increased OXPHOS and ATP production. This may be attributed to *A. castellanii* boosting ATP production to enhance cellular processes involved in combating the pathogen. Conversely, alterations to mitochondria dynamics induced by *C. jejuni* infection could have influenced ETC activity, however, this remains to be elucidated.

Mitochondria are also the main source of cellular reactive oxygen species (ROS) [48]. Despite observing a ∼12-fold increase in ROS levels at 1 hour post-infection, it was surprising that ROS returned to levels similar to those of uninfected cells by 3 hrs post-infection. This observation prompted us to reconsider the purpose of mitochondria recruitment to the *C. jejuni*-containing vacuole. While traditionally viewed as a mechanism for antimicrobial activity [49], our findings suggest an alternative hypothesis. We propose that the recruitment of mitochondria to the *C. jejuni*-containing vacuole may serve a different, yet unknown function, potentially manipulated by the bacteria themselves. This manipulation of mitochondria dynamics and their recruitment can provide *C. jejuni* with advantages such as access to nutrients [50]. *C. jejuni* exhibits distinctive metabolic characteristics, setting it apart from other enteropathogenic bacteria, notably its limited carbohydrate catabolism. Indeed, the mitochondria serve as a hub for amino acid synthesis, including the production of glutamine, glutamate, proline, and aspartate [51]. All are capable of supporting *C. jejuni* viability [33] and growth. Previously, we reported significant upregulation of the Peb1 system in intra-amoebae *C. jejuni*, an ABC transporter responsible for the uptake of aspartate and glutamate [16]. Mutation in *peb1c* (*cj0922c*), an ATP-binding protein responsible for energy coupling to the peb1ABC transporter, did not exhibit a survival defect **(Supplementary File 3 (Figure S5))**, suggesting potential redundancy in the system [33].

In addition to amino acid synthesis and ATP production, mitochondria play a crucial role in regulating cellular calcium homeostasis [52], which is essential for various cellular functions. Mitochondria act as buffers, sequestering and releasing calcium ions in response to intracellular signals [52]. The increase in cytosolic calcium levels in *A. castellanii* triggered by *C. jejuni*, as demonstrated in previous studies using mammalian cells [53], leads to speculation regarding potential connections between calcium homeostasis and the observed alterations in mitochondria during *C. jejuni* infection.

Various studies indicate that bacteria trigger distinct metabolic programs in phagocytes during infection, and this metabolic rewiring is thought to be crucial for bacterial replication and/or survival, as well as host response to the infection [32, 54]. Here, we present evidence that supports *A. castellanii* rewires its metabolic activities at early stages of the infection. Interestingly, these metabolic activities return to levels comparable to those of uninfected cells as the infection progresses. We noted upregulation of phosphoenolpyruvate carboxykinases (PEPCK), responsible for converting oxaloacetate to phosphoenolpyruvate (PEP), a precursor of glucose [55]. PEPCK, coupled with enhanced glucose-6-phosphate (G6P) and increased lactate production suggested a metabolic shift towards glycolysis [56]. This metabolic adaptation may be driven by the demand for glucose as an energy source to counter *C. jejuni* infection. These findings are consistent with a metabolic response resembling the “Warburg effect” phenomenon, which is commonly observed in cancer cells and often associated with increased aerobic glycolysis [32]. Aerobic glycolysis generates less ATP per molecule of glucose compared to OXPHOS but allows for the rapid production of ATP and other essential biomolecules needed to mount an effective response to the infection [57].

Previously, it was suggested that *A. castellanii* might promote the survival and proliferation of extracellular *C. jejuni* through mechanism of amoeba-mediated depletion of dissolved oxygen [34]. Presently, we propose that amoeba lactate production may also play additional and an important role in this phenomenon. The increased aerobic glycolysis produced lactate as by-product, our data revealed a rapid depletion of extracellular lactate in the presence of *C. jejuni*. This aligns with our previous study, in which we reported a significant upregulation of LctP, an l-lactate permease [16]. Therefore, we propose that *C. jejuni* effectively exploits lactate produced by *A. castellanii* as an additional energy source which contributes to its survival within the host and in shared environmental niches [17, 35, 58]. This hypothesis also supports recent findings that showed lactate transport regulation is increased in aerobic environments [17], thereby reinforcing the notion that *C. jejuni* effectively utilizes lactate produced by *A. castellanii* as an additional energy source, contributing to its survival in shared environmental niches.

In conclusion, this study represents a significant advancement in our understanding of the interaction between *C. jejuni* and *A. castellanii* and provide general insight into host pathogen interactions. It provides evidence that *C. jejuni* manipulates its transient amoeba host to facilitate its survival. The findings enlighten us on the sophisticated strategies employed by *C. jejuni* to persist within host cells and the complex dynamics of microbial pathogenesis. Moreover, this study suggests that the *A. castellanii* model could be utilized to study *C. jejuni* host-pathogen dynamics, offering promising opportunities for further understanding and potentially mitigating the impact of *C. jejuni* infections.

## Methods

### Strains and Cultures

*Campylobacter jejuni* strain 11168H, a motile derivative of the original sequence strain NCTC 11168 [59], were stored using Protect bacterial preservers (Technical Service Consultants, England, UK) at −80 °C as previously described. Bacteria were streaked on Columbia blood agar (CBA) plates containing Columbia agar base (Oxoid) supplemented with 7% (v/v) horse blood (TCS Microbiology, UK) and grown at 37 °C in a microaerobic chamber (Don Whitley Scientific, England, UK), containing 85% N2, 10% CO_2_, and 5% O_2_for 48 h. Bacteria were grown on CBA plates for a further 16 h at 37 °C prior to use.

*Acanthamoeba castellanii* acquired from Culture Collection of Algae and Protozoa (CCAP) 1501/10 (Scottish Marine Institute) were grown to confluence at 25 °C in 75 cm^2^ tissue culture flasks containing peptone yeast and glucose (PYG) media (proteose peptone 20 g, glucose 18 g, yeast extract 2 g, sodium citrate dihydrate 1 g, MgSO4 × 7H_2_O 0.98 g, Na_2_HPO4 × 7H_2_O 0.355 g, KH_2_PO4 0.34 g in distilled water to make 1000 mL, pH was adjusted to 6.5). Amoebae were harvested into suspension, and viability was determined by counting using inverted light microscope.

T84 (human intestinal cell line) were grown in Dulbecco’s modified Eagle’s medium and Ham’s F-12 (DMEM/F-12) supplemented with 10L% FBS and 1L% non-essential amino acid. The monolayers, ∼10^5^, were seeded in a 24-well tissue culture plates and were grown up to ∼10^6^ in a 5L% CO2 atmosphere and were then infected with C. jejuni 11168H at an m.o.i. of 200L:L1 for 4 hrs including 1 hr of mitochondrial staining (mitochondrial staining is described below in detail).

### *C. jejuni* mutant construction

Isothermal assembly (ISA) cloning, based on the method described by Gibson et al. (58), was used to generate constructs for *C. jejuni* mutagenesis. Genes of interest were disrupted by inserting antibiotic resistance cassette (encoding kanamycin resistance (Kan^r^)) or Apramycin (Ap^r^) into the reading frame. The plasmids generated contain upstream and downstream gene flanks which are recombined by double homologous crossover into the *C. jejuni* chromosome after electroporation and antibiotic selection. HiFi DNA assembly cloning kit (New England Biolabs (NEB)) was used according to manufacturer’s instruction. Briefly, *C. jejuni* 11168H wildtype was used to generate a Δ*cdtABC* (Ap^r^) and Δ*lctP* (Kan^r^) mutants, *C. jejuni* ΔcjeN (*cj0979c*) mutant was described previously [16]. Four fragments for HiFi DNA assembly were prepared as follows, pGEM3Zf^(–)^ (Promega) was digested with HincII and phosphatase-treated. Kan^r^ resistance cassette was PCR-amplified primers **(Table S3)**. Primers were designed to amplify flanking regions of the gene to be deleted, removing most of the open reading frame. The left flanking region was termed fragment 1 (F1; beginning of gene of interest plus upstream flanking DNA) and the right flanking region fragment 2 (F2; end of gene of interest plus downstream flanking DNA (Supplementary file 3 (**Table S3)**). All PCRs were performed using Q5 high-fidelity DNA polymerase (NEB). Plasmids with desired mutation were cloned in *Escherichia coli* DH5α, and transformants were selected on LB agar supplemented with apramycin and ampicillin. The resultant plasmids were electroporated into *C. jejuni* with corresponding antibiotic selection. Colonies were screened by PCR to confirm the correct insertion of the cassette into the genome.

Complementation of Δ*cjeN*, Δ*lctp* and Δ*ciaB* were generated using the plasmid pRRA (A=Apramycin) as previously described [37, 60]. Briefly, the wild type *cjeN*, *lctP* and *ciaB* genes with intact native promoter was amplified and the fragments was inserted into plasmid pRRA [Apramycin resistant (Ap^r^)]; this plasmid introduces genes into the conserved multi-copy rRNA gene clusters on the chromosome. Plasmids were cloned in *E. coli* DH5α, and transformants were selected on LB agar supplemented with Ap^r^. The resultant plasmids were electroporated into *C. jejuni* with selection for Kan^r^ and Ap^r^ Supplementary file 3 (**Table S3)**.

### *C. jejuni* recombinant protein expression and purification

Recombinant CjeN_his6_ (Cj0979c) and its variant CjeN_Thr69Ser77Thr111his6_ were cloned into pET21a^(+)^ plasmid using a NEBuilder HiFi DNA assembly cloning kit (New England Biolabs). The proteins were overexpressed in *E. coli* strain BL21(DE3). The cells were grown in LB broth containing 150 μg/ml ampicillin at 37°C overnight. Cells were harvested by centrifugation at 4,000L×Lg for 15 min at 4°C and resuspended in lysis buffer; 20LmM Tris-HCl, pH 8.0, 2500LmM NaCl, 5% [vol/vol] glycerol, containing EDTA-free cOmplete protease inhibitor mixture (Roche) and 15 mM imidazole. Cell debris was removed by centrifugation at 10,000L×Lg for 30 min at 4°C. The supernatant was incubated with Ni-NTA agarose nickel-charged resins (Qiagen) that had been equilibrated in lysis buffer. The protein-bound resin was washed with lysis buffer containing 400 mM imidazole to elute recombinant protein. The primers used to generate recombinant proteins are presented in Supplementary file 3 (**Table S3)**.

### Differential gene expression analysis

*A. castellanii* infected with *C. jejuni* strain 11168H was defined as the ‘test group’ and uninfected *A. castellanii* as the control (incubated in the absence of bacteria) after a total of 4 h. Experiments were performed as described above; briefly, *A. castellanii* 10^6^ were infected with *C. jejuni* 11168H (M.O.I of 200) for 3 h and washed three times to remove extracellular bacteria. Gentamycin was added to a final concentration of 100 μg/mL; this culture was incubated for a further 1 h before washing three times and RNA was extracted using Triazole (Sigma Aldrich, England, UK) following manufacturer’s protocol. RNA was sent to Genewiz UK (Azenta) for processing and paired-end RNA sequencing (Illumina).

Illumina 150 bp Paired-end reads were trimmed using trimmomatic V 0.39; with parameters leading: 3, trailing 3 and MINLEN: 36, TruSeq3-PE adapter removal was also conducted. Abundance estimates were quantified using Kallisto (v0.46.1) with default parameters against reference sequence, Acastellanii.strNEFF v1 GCA_000313135.1. Differential gene expression analysis was conducted using edgeR (empirical analysis of DGE in R) in R with adjusted p-value cut-off of <0.05, and log fold change cut-off of >2.

For Gene ontology (GO) annotation and enrichment analysis; *A. castellanii* Neff strain was first De Novo assembled using Trinity v2.15.1 assembler. Followed on by quantification using RSEM (RNA-Seq by Expectation-Maximization) [61] and differential expression analysis by edgeR on OmicsBox (Biobam). GO mapping was done using OmicsBox (biobam).

### Gene expression by Real-Time RT-qPCR

Expression of genes of interest were quantified by real-time RT-qPCR and normalized against *A. castellanii* 18s. A 1 μg of total RNA of each sample was reverse-transcribed to cDNA using RT2 first strand kit (Qiagen) according to manufacturer’s protocol. Quantification of gene expression was achieved by real-time RT-qPCR using Sybr^TM^ Green real-time PCR master mix using primers generated using Primer Quest (IDT) **(Supplementary File S1 (Table S4))**. Real-time RT-PCR was performed in 96-well plates using an ABI PRISM 7300 Real-time PCR System (Applied Biosystems), and the relative gene expression for the different genes was calculated from the crossing threshold (Ct) value according to the manufacturer’s protocol (2^−ΔΔCt^) after normalization using the 18s endogenous control [62].

### Survival Assay

Invasion and survival assays were performed as previously described [15]. Briefly, *C. jejuni* 11168H and its mutants were incubated with a monolayer of approximately 10^6^ A*. castellanii* at a multiplicity of infection (M.O.I) of 100:1 for 3 h at 25 °C in 2 mL PYG media (the M.O.I. of 100:1 was chosen to allow maximal internalization of bacteria without impacting amoebae predation through density-dependent inhibition). The monolayer was washed 3× with phosphate buffer saline solution (PBS) and incubated for 1 h in 2 mL of PYG media containing 100 μg/mL of gentamicin.

*C. jejuni* cells were harvested by scraping the amoebae into suspension and centrifuged for 10 min at 350× g to pellet the bacteria and amoebae. Supernatant was discarded and the pellet was suspended in 1 mL of distilled water containing 0.1% (v/v) Triton X-100 for 10 min at room temperature with vigorous pipetting (every 2 min) to lyse the amoebae and release bacterial cells. The suspension was then centrifuged for a further 10 min at 4000× g, the resultant pellet was resuspended in 1 mL PBS and enumerated for colony forming units on CBA plates for up to 72 h at 37 °C microaerobically.

### DNA double stranded breakages and Mitochondria imaging

*A. castellanii* cells were infected with *C. jejuni* 11168H wildtype and its mutants as described above. After infection, amoeba cells were washed to remove residual media and then fixed with 4% paraformaldehyde for 20 minutes. Subsequently, the cells were permeabilized with 0.1% Triton-X100 in phosphate-buffered saline (PBS) for 5 minutes, followed by additional washes with PBS. Samples were mounted using Fluoroshield™ with DAPI before imaging with an inverted confocal microscope.

To assess double-stranded DNA breakages, *A. castellanii* cells were infected with *C. jejuni* 11168H wildtype strain, its Δ*cdtABC*, Δ*cjeN* (*cj0979c*) mutants, or CjeN_his6_-coupled latex beads for a total of 4 hours. CjeN_his6_-coupled latex beads were prepared according to the manufacturer’s protocol. Briefly, plain latex beads were suspended in borate buffer, washed, and then coupled with CjeN_his6_ protein. Protein quantification was performed using Pierce™ BCA Protein Assay Kit. Following coupling, beads were blocked with 5% BSA in borate buffer and washed before resuspension in PBS (pH 7.4). *A. castellanii* cells were then incubated with 10 μL (∼0.3ng of total protein), CjeN_his6_-coupled beads for 3 hrs. *A. castellanii* cells were subsequently washed with PBS, fixed, permeabilized, and incubated with Rabbit anti-γH2AX antibody (Abcam, UK) followed by Alexa Fluor® 488 Goat anti-Rabbit IgG Cross-Adsorbed Secondary staining.

All CjeN *in vitro* nuclease activity assays were performed in Tris-HCl (pH 7.5) buffer containing 130 µM calcium (Ca^2+^), 5 mM magnesium (Mg^2+^) and 5% glycerol at 37 °C for 30 mins and samples were ran on 1% Agarose gel. Mitochondria imaging was performed as follows, at specified intervals following infection *A. castellanii* were incubated with Mitobrilliant^TM^ 646 for 1 hr. Cells were washed and fixed before permeabilization and imaging.

### Imaging and Microscopy

Experiments were performed in 35 mm μ-Dish devices (IBIDI) unless otherwise stated. Confocal laser scanning microscopy and super-resolution Airyscan images were obtained using an inverted Zeiss LSM 880 confocal microscope (Zeiss, Berlin, Germany). Images were taken at 5 s intervals for time lapse experiments. Alexa fluor® 488 was monitored in the green region (excitation (EX) and emission (EM) wavelengths are 488/510 nm). Mitobrilliant^TM^ 646 was monitored in the red region (EX/EM= 655/668 nm). 4′,6-diamidino-2-phenylindole (DAPI) was monitored in the blue region (EX/EM=355/405).

### Calcium and Reactive oxygen species (ROS) quantification assay

Calcium dysregulation was monitored using Fluo-4 NW Calcium Assay Kit (Thermofischer scientific, US) according to manufacturer’s instructions. Briefly, *A. castellanii* were infected with *C. jejuni* 11168H in a 96 well plate as described above. Cells were washed 3 x with PBS and 100 µL of the dye loading solution was added to each sample well. Plate was incubated at 25°C for 1 h prior to reading using SpectraMax iD5 (Molecular Devices). Fluo-4 NW Calcium Assay Kit (excitation 494 nm and emission at 516 nm).

ROS was quantified using CellROX™ Deep Red (Thermofischer) following manufactures instructions. Briefly, after infection *A. castellanii* cell were washed and 5 μM of CellROX Red reagent was added to the cells for 30 mins. Cells were washed three times with PBS before analysing with SpectraMax iD5. CellROX™ Deep Red (excitation 640 nm and emission at 665 nm)

### Lactate and Glucose-6-phosphate assay

To determine lactate and Glucose-6-phosphate (G6P) concentrations in *A. castellanii* infected with *C. jejuni*, cells were infected as described above for 2 hrs, 4 hrs and 24 hrs. Following infection, cells were homogenized and subjected to deproteinization using a Deproteinizing Sample Preparation Kit (Merk) following the manufacturer’s instructions. The samples were analysed for lactate using Lactate-Glo^TM^ Assay Kit (Promega) and Amplite® Fluorimetric Glucose-6-Phosphate Assay Kit (AAT Bioquest) was used to measure G6P concentrations according to the manufacturer’s instructions with a SpectraMax iD5. Lactate standards provided in the kits were used to determine absolute concentrations.

### Statistical Analysis

All experiments presented are at least three independent biological replicates. RNA-Seq analysis were performed in R using the combined data generated from the bioinformatics. Evaluation of *A. castellanii* interaction with *C. jejuni* was based on at least 200 amoebae cells per experiment. Differentially expressed genes were considered significant when the *p*-value of three independent biological experiments was below <0.05. All other data were analysed using Prism statistical software (Version 9, GraphPad Software, San Diego, CA, USA), and statistical significance was considered when the *p*-value was <0.05. All values are presented as standard deviation.

## Supplementary Material

Supplementary File 1: All differential regulated genes in *A. castellanii* infected with *C. jejuni*; Supplementary File 2: All GO Term enrichment genes in *A. castellanii* infected with *C. jejuni*; Figure S1: Proteosome 20s activity assay; Figure S2: Multiple Sequence Alignment of *C. jejuni* CjeN, *S. aureus* Nuc1 and *B. subtilis* YncB protein sequences.; Figure S3: CiaB mutant is trafficked to digestive vacuoles at a faster rate than the wildtype and is not associated with host mitochondria. Table S2: Table of all the strain used in this study; Table S3: Primers used to make mutants and recombinant proteins; Table S4: RT-qpCR primers.

## Author Contributions

F.N conceived the study. F.N. designed the experiments. F.N. and BL. performed experiments, RNA-Seq data analysis. F.N wrote the manuscript. F.N, B.W.W, B.L, and R.A.S edited the manuscript. All authors have read and agreed to the manuscript.

## Data availability statement

The data that supports the RNA-Seq findings of this study are available in the Gene Expression Omnibus (GEO) under data set identifier GSE262802.

## Reference

1. Berlanga, M., M. Viñas, and R. Guerrero, Prokaryotic Basis of Eukaryotic Eco-Evo Development. Developmental Biology in Prokaryotes and Lower Eukaryotes, 2021: p. 313–330.

2. Pallen, M.J. and B.W. Wren, Bacterial pathogenomics. Nature, 2007. 449(7164): p. 835–42.

3. Khan, N.A. and R. Siddiqui, Predator vs aliens: bacteria interactions with Acanthamoeba. Parasitology, 2014. 141(7): p. 869–74.

4. Henriquez, F.L., et al., Paradigms of Protist/Bacteria Symbioses Affecting Human Health: Acanthamoeba species and Trichomonas vaginalis. Front Microbiol, 2020. 11: p. 616213.

5. Siddiqui, R. and N.A. Khan, Biology and pathogenesis of Acanthamoeba. Parasit Vectors, 2012. 5(1): p. 6.

6. Shi, Y., et al., The Ecology and Evolution of Amoeba-Bacterium Interactions. Appl Environ Microbiol, 2021. 87(2).

7. Mungroo, M.R., R. Siddiqui, and N.A. Khan, War of the microbial world: Acanthamoeba spp. interactions with microorganisms. Folia Microbiol (Praha), 2021. 66(5): p. 689–699.

8. Elmi, A., et al., Revisiting Campylobacter jejuni Virulence and Fitness Factors: Role in Sensing, Adapting, and Competing. Front Cell Infect Microbiol, 2020. 10: p. 607704.

9. Kaakoush, N.O., et al., Global Epidemiology of Campylobacter Infection. Clin Microbiol Rev, 2015. 28(3): p. 687–720.

10. Lehri, B., et al., Phenotypic and genotypic characterization of Campylobacter coli isolates from the Vietnamese poultry production network; a pilot study. Frontiers in Industrial Microbiology, 2024. 2.

11. Jones, K., The Campylobacter conundrum. Trends Microbiol, 2001. 9(8): p. 365–6.

12. Bare, J., et al., Influence of temperature, oxygen and bacterial strain identity on the association of Campylobacter jejuni with Acanthamoeba castellanii. FEMS Microbiol Ecol, 2010. 74(2): p. 371–81.

13. Axelsson-Olsson, D., et al., Protozoan Acanthamoeba polyphaga as a potential reservoir for Campylobacter jejuni. Appl Environ Microbiol, 2005. 71(2): p. 987–92.

14. Vieira, A., A.M. Seddon, and A.V. Karlyshev, Campylobacter-Acanthamoeba interactions. Microbiology (Reading), 2015. 161(Pt 5): p. 933–947.

15. Nasher, F. and B.W. Wren, Transient internalization of Campylobacter jejuni in Amoebae enhances subsequent invasion of human cells. Microbiology (Reading), 2022. 168(2).

16. Nasher, F., et al., Survival of Campylobacter jejuni 11168H in Acanthamoebae castellanii Provides Mechanistic Insight into Host Pathogen Interactions. Microorganisms, 2022. 10(10).

17. Sinha, R., et al., Gut metabolite L-lactate supports Campylobacter jejuni population expansion during acute infection. Proc Natl Acad Sci U S A, 2024. 121(2): p. e2316540120.

18. Wang, N., et al., Ac-HSP20 Is Associated With the Infectivity and Encystation of Acanthamoeba castellanii. Front Microbiol, 2020. 11: p. 595080.

19. Cheeseman, I.M. and A. Desai, Molecular architecture of the kinetochore-microtubule interface. Nat Rev Mol Cell Biol, 2008. 9(1): p. 33–46.

20. Luo, D., et al., Universal Stress Proteins: From Gene to Function. Int J Mol Sci, 2023. 24(5).

21. Palmieri, F. and C.L. Pierri, Mitochondrial metabolite transport. Essays Biochem, 2010. 47: p. 37–52.

22. Hardie, D.G., AMP-activated/SNF1 protein kinases: conserved guardians of cellular energy. Nat Rev Mol Cell Biol, 2007. 8(10): p. 774–85.

23. Pensalfini, A., et al., Assessing Rab5 Activation in Health and Disease. Methods Mol Biol, 2021. 2293: p. 273–294.

24. Ge, S.X., D. Jung, and R. Yao, ShinyGO: a graphical gene-set enrichment tool for animals and plants. Bioinformatics, 2020. 36(8): p. 2628–2629.

25. Lara-Tejero, M. and J.E. Galan, A bacterial toxin that controls cell cycle progression as a deoxyribonuclease I-like protein. Science, 2000. 290(5490): p. 354–7.

26. Rangarajan, E.S. and V. Shankar, Sugar non-specific endonucleases. FEMS Microbiol Rev, 2001. 25(5): p. 583–613.

27. Almagro Armenteros, J.J., et al., SignalP 5.0 improves signal peptide predictions using deep neural networks. Nature Biotechnology, 2019. 37(4): p. 420–423.

28. Torriglia, A., et al., On the use of Zn2+ to discriminate endonucleases activated during apoptosis. Biochimie, 1997. 79(7): p. 435–8.

29. Siddiqui, R. and N.A. Khan, Biology and pathogenesis of Acanthamoeba. Parasit Vectors, 2012. 5: p. 6.

30. Shekhova, E., Mitochondrial reactive oxygen species as major effectors of antimicrobial immunity. PLoS Pathog, 2020. 16(5): p. e1008470.

31. Rossi, A., P. Pizzo, and R. Filadi, Calcium, mitochondria and cell metabolism: A functional triangle in bioenergetics. Biochim Biophys Acta Mol Cell Res, 2019. 1866(7): p. 1068–1078.

32. Escoll, P. and C. Buchrieser, Metabolic reprogramming of host cells upon bacterial infection: Why shift to a Warburg-like metabolism? FEBS J, 2018. 285(12): p. 2146–2160.

33. Stahl, M., J. Butcher, and A. Stintzi, Nutrient acquisition and metabolism by Campylobacter jejuni. Front Cell Infect Microbiol, 2012. 2: p. 5.

34. Bui, X.T., et al., Survival of Campylobacter jejuni in co-culture with Acanthamoeba castellanii: role of amoeba-mediated depletion of dissolved oxygen. Environ Microbiol, 2012. 14(8): p. 2034–47.

35. Thomas, M.T., et al., Two respiratory enzyme systems in Campylobacter jejuni NCTC 11168 contribute to growth on L-lactate. Environ Microbiol, 2011. 13(1): p. 48–61.

36. Patrick, M.E., et al., Features of illnesses caused by five species of Campylobacter, Foodborne Diseases Active Surveillance Network (FoodNet) - 2010-2015. Epidemiol Infect, 2018. 146(1): p. 1–10.

37. Nasher, F. and B.W. Wren, Flagellin O-linked glycans are required for the interactions between Campylobacter jejuni and Acanthamoebae castellanii. Microbiology (Reading), 2023. 169(8).

38. Mehta, A. and J.E. Haber, Sources of DNA double-strand breaks and models of recombinational DNA repair. Cold Spring Harb Perspect Biol, 2014. 6(9): p. a016428.

39. Cannan, W.J. and D.S. Pederson, Mechanisms and Consequences of Double-Strand DNA Break Formation in Chromatin. J Cell Physiol, 2016. 231(1): p. 3–14.

40. Ammendola, S. and A. Battistoni, New Insights into the Role of Metals in Host-Pathogen Interactions. Int J Mol Sci, 2022. 23(12).

41. Beiter, K., et al., An endonuclease allows Streptococcus pneumoniae to escape from neutrophil extracellular traps. Curr Biol, 2006. 16(4): p. 401–7.

42. Ramesh, A., Investigation of Molecular Mechanisms of Biofilm Dispersal and Attempts to Create an Inducible Gene Expression System in Campylobacter. 2020: Kingston University.

43. Rada, B., Neutrophil Extracellular Traps. Methods Mol Biol, 2019. 1982: p. 517–528.

44. Newsome, A.L., et al., Interactions between Naegleria-Fowleri and Legionella-Pneumophila. Infection and Immunity, 1985. 50(2): p. 449–452.

45. Abarca-Rojano, E., et al., Mycobacterium tuberculosis virulence correlates with mitochondrial cytochrome c release in infected macrophages. Scand J Immunol, 2003. 58(4): p. 419–27.

46. Fischer, S.F., et al., Protection against CD95-induced apoptosis by chlamydial infection at a mitochondrial step. Infection and Immunity, 2004. 72(2): p. 1107–1115.

47. Dorn, G.W., 2nd, Evolving Concepts of Mitochondrial Dynamics. Annu Rev Physiol, 2019. 81: p. 1–17.

48. Turrens, J.F., Mitochondrial formation of reactive oxygen species. J Physiol, 2003. 552(Pt 2): p. 335–44.

49. Holmbeck, M.A. and G.S. Shadel, Mitochondria provide a ‘complex’ solution to a bacterial problem. Nat Immunol, 2016. 17(9): p. 1009–10.

50. Spier, A., F. Stavru, and P. Cossart, Interaction between Intracellular Bacterial Pathogens and Host Cell Mitochondria. Microbiol Spectr, 2019. 7(2): p. 10.1128/microbiolspec.bai-0016-2019.

51. Li, Q. and T. Hoppe, Role of amino acid metabolism in mitochondrial homeostasis. Front Cell Dev Biol, 2023. 11: p. 1127618.

52. Matuz-Mares, D., et al., Mitochondrial Calcium: Effects of Its Imbalance in Disease. Antioxidants (Basel), 2022. 11(5): p. 801.

53. Hu, L., R.B. Raybourne, and D.J. Kopecko, Ca2+ release from host intracellular stores and related signal transduction during Campylobacter jejuni 81-176 internalization into human intestinal cells. Microbiology (Reading), 2005. 151(Pt 9): p. 3097–3105.

54. Price, C., et al., Paradoxical Pro-inflammatory Responses by Human Macrophages to an Amoebae Host-Adapted Legionella Effector. Cell Host Microbe, 2020. 27(4): p. 571–584 e7.

55. Yang, J., S.C. Kalhan, and R.W. Hanson, What is the metabolic role of phosphoenolpyruvate carboxykinase? J Biol Chem, 2009. 284(40): p. 27025–9.

56. Chaudhry, R. and M. Varacallo, Biochemistry, glycolysis. 2018.

57. Soto-Heredero, G., et al., Glycolysis - a key player in the inflammatory response. FEBS J, 2020. 287(16): p. 3350–3369.

58. Hofreuter, D., Defining the metabolic requirements for the growth and colonization capacity of Campylobacter jejuni. Front Cell Infect Microbiol, 2014. 4: p. 137.

59. Karlyshev, A.V., et al., A novel paralogous gene family involved in phase-variable flagella-mediated motility in Campylobacter jejuni. Microbiology (Reading), 2002. 148(Pt 2): p. 473–480.

60. Karlyshev, A.V. and B.W. Wren, Development and application of an insertional system for gene delivery and expression in Campylobacter jejuni. Appl Environ Microbiol, 2005. 71(7): p. 4004–13.

61. Langmead, B. and S.L. Salzberg, Fast gapped-read alignment with Bowtie 2. Nat Methods, 2012. 9(4): p. 357–9.

62. Nachamkin, I., X.H. Yang, and N.J. Stern, Role of Campylobacter jejuni flagella as colonization factors for three-day-old chicks: analysis with flagellar mutants. Appl Environ Microbiol, 1993. 59(5): p. 1269–73.

