## Supplementary File 3 for "*Acanthamoeba castellanii* as a model for unveiling *Campylobacter jejuni* host-pathogen dynamics"

***Acanthamoeba castellanii* as a model for unveiling *Campylobacter jejuni* host-pathogen dynamics.**

Fauzy Nasher^1*^, Burhan Lehri^1^, Richard Stabler^1^, Brendan W. Wren^1*^.

^1^Department of Infection Biology

London School of Hygiene and Tropical Medicine

Keppel St, London WC1E 7HT

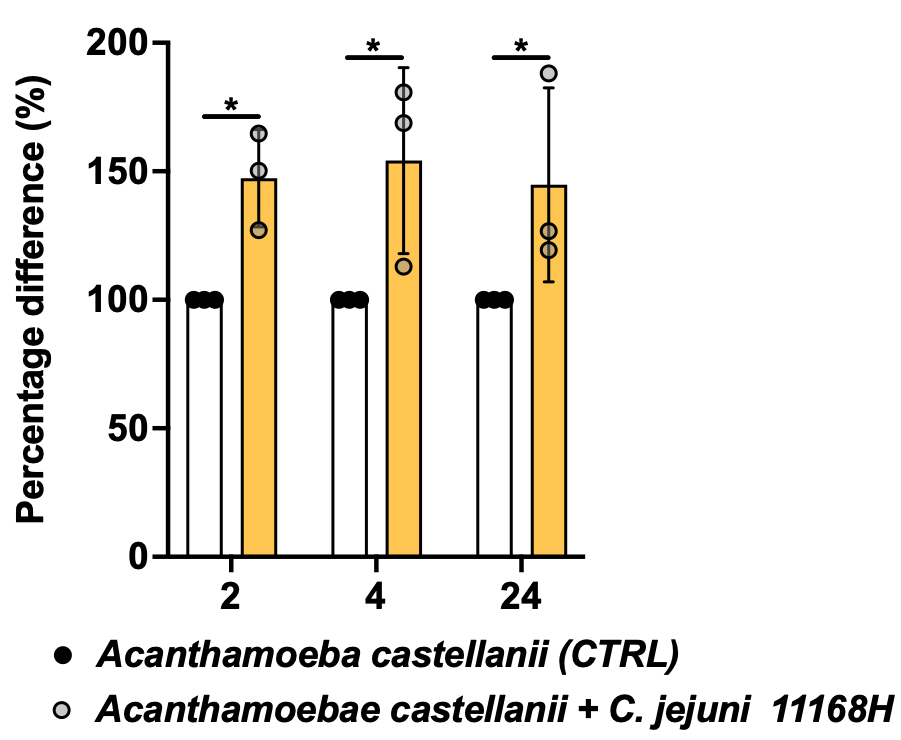

**Figure S1. Proteosome 20s activity assay.** Proteosome (20S) activity was monitored using Amplite® Fluorimetric Proteasome 20S Activity Assay Kit Green Fluorescence (AAT Bioquest) according to manufacturer’s instructions. Amplite® Fluorometric 20S Proteasome Assay Kit uses LLVY-R110 as a fluorogenic indicator for proteasome activity. Briefly, amoeba cells were infected with *C. jejuni* 11168H in a 96 well plate as described above, cells were washed 3x with PBS and 100 µL proteasome working solution was added per well. The plate was incubated at 25°C for 2 h and fluorescence intensity was monitored at Ex/Em = 490/525 nm using SpectraMax iD5. Significance levels are indicated as follows: *p<0.05; error bars represent standard deviation (SD).

**Multiple Sequence Alignment**

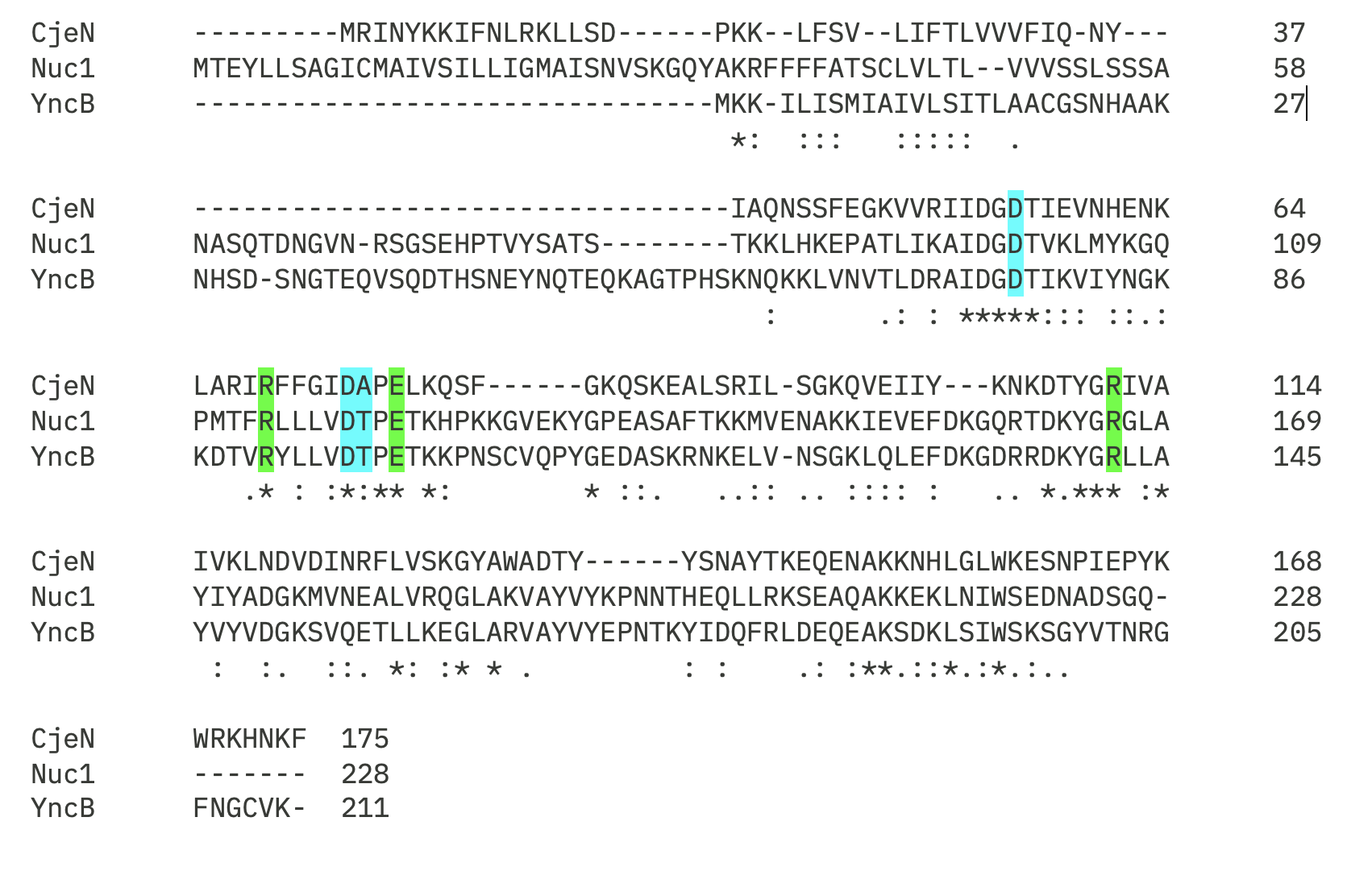

**Figure S2. Multiple Sequence Alignment of *C. jejuni* CjeN, *S. aureus* Nuc1 and *B. subtilis* YncB protein sequences.** Green indicates conserved active sites Arg 69 and Glu77 and Argi 111 while blue indicate metal binding sites, Asp 55, 74 and Ala 75. Clustal omega was used for alignment.

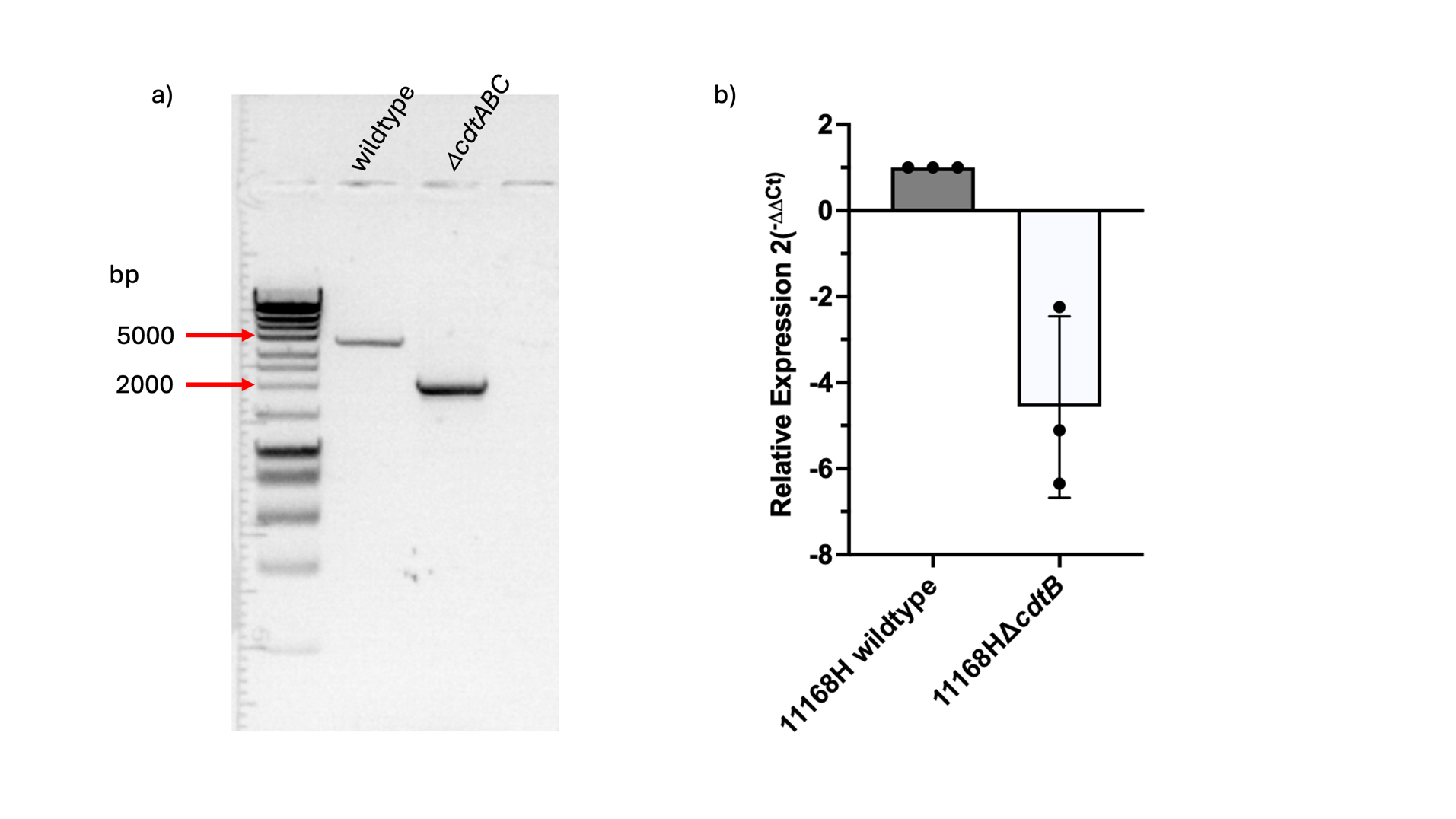

**Figure S3: *C. jejuni* 11168HΔ*cdtABC* mutant. a)** Agarose gel (1%) electrophoresis showing PCR products of 11168H wildtype and its Δ*cdtABC* mutant (HyperLadder^TM^ 1kb); **b)** RT-qPCR showing expression levels of Δ*cdtABC* mutant relative to the wildtype after normalization with *gyrA* endogenous control.

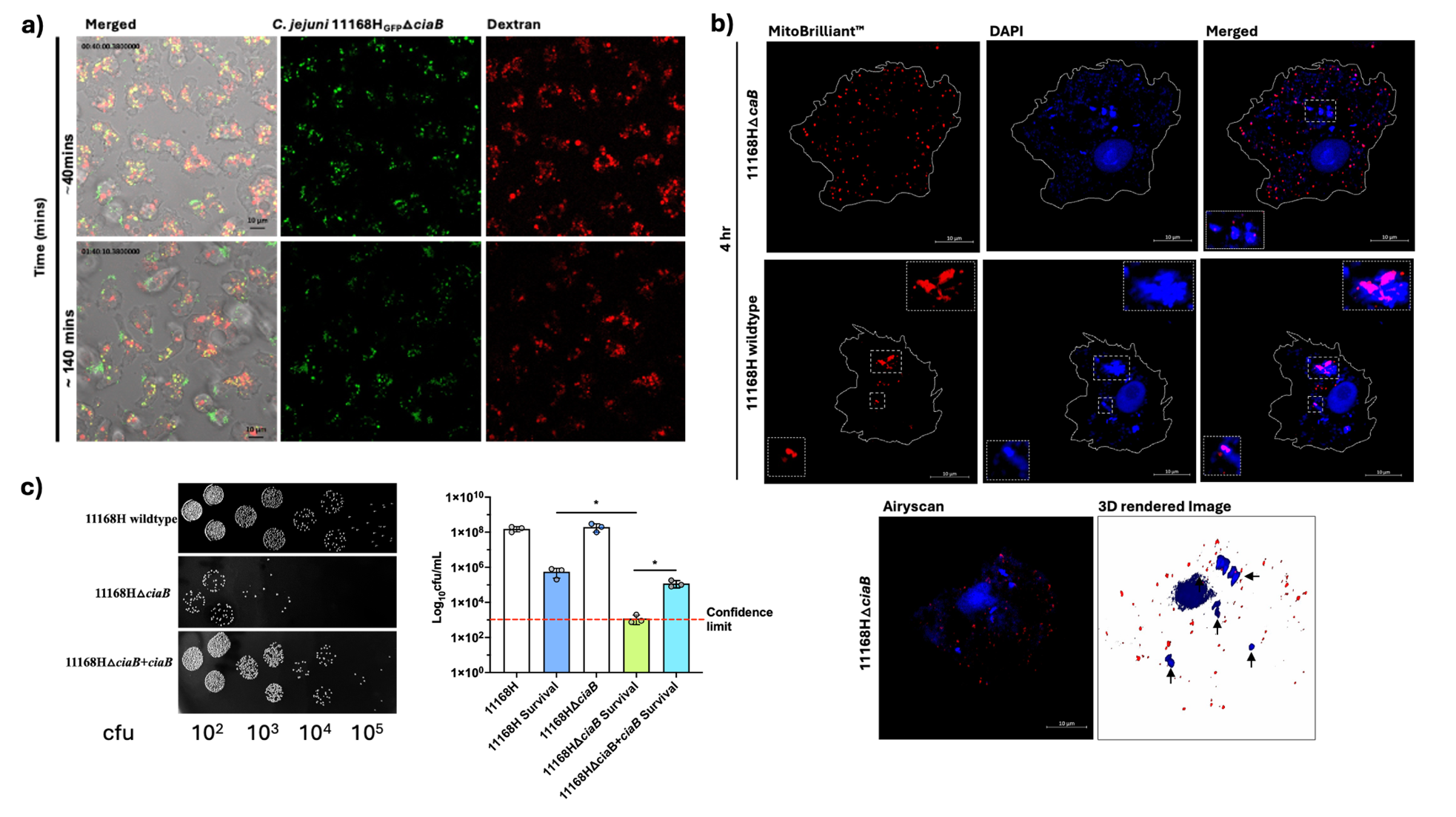

**Figure S4: CiaB mutant is trafficked to digestive vacuoles at a faster rate than the wildtype and ciaB mutant is not associated with host mitochondria. a)** Time lapse imaging showing *C. jejuni* 11168HΔ*ciaB* mutant ~40 mins and 1 hr 40 mins post infections; interestingly ciaB mutants are trafficked to digestive vacuole at a faster rate than what we usually observe [1]. **b)** CiaB mutant was not associated with host mitochondria and **c)** colony forming units (cfu) of *C. jejuni* 11168H wildtype relative to its Δ*ciaB* and Δ*ciaB*+*ciaB* mutant after 4 hr infection (including 1 hr gentamycin treatment), wildtype strain showed 2-fold survival higher than the Δ*ciaB* mutant strain. Mutants were constructed as described in the methods section. Arrows indicate bacteria. Error bars are presented at standard deviation (SD); **p*<0.05.

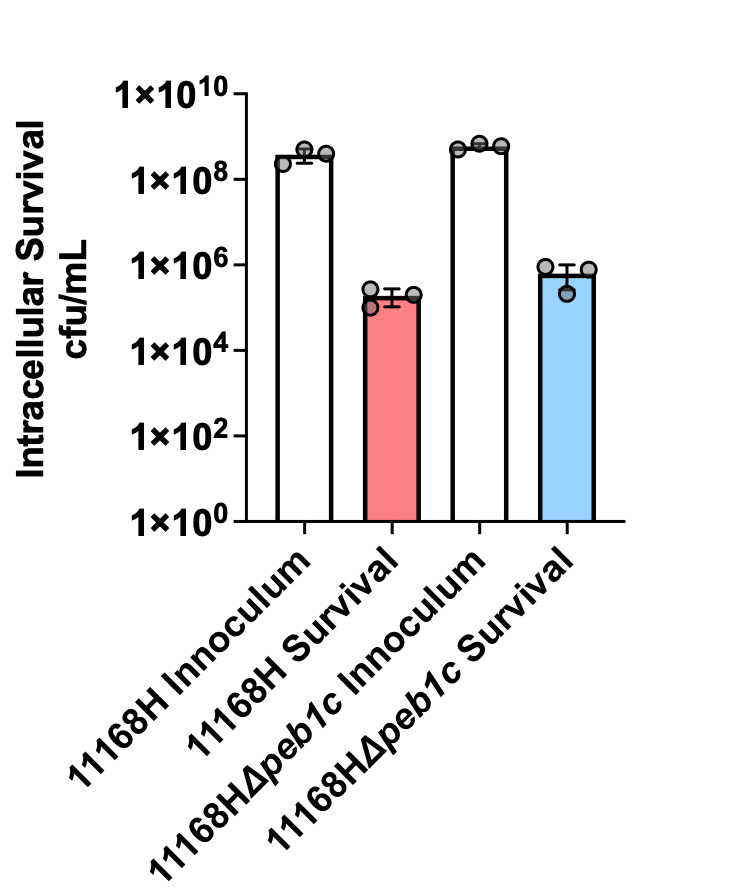

**Figure S5: *peb1c* mutant intracellular survival was similar to that of the wildtype strain.** *C. jejuni* 11168H Δ*pe1bC* mutant construct was obtained from was acquired from obtained from the Campylobacter Resource Facility (<http://crf.lshtm.ac.uk/wren_mutants.htm>, accessed on 14 April 2024) and strain 11168H was naturally transformed using the previously described biphasic method [2]. Survival assay was performed as described in the methods.

**Table S3: Primers used to make mutants and recombinant proteins.**

| **Primer Name** | **Sequence** | |
| --- | --- | --- |
| **Mutagenesis Primers** | | |
| cdtABC_F1 | **Fwd:** 5-ccggggatcctctagagtcggcggaaaattataatgaaattta-3 | **Rvs:** 5- gcccaaaccgttaaagctgc -3 |
| cdtABC_Apr | **Fwd:** 5- cggtttgggcgtaacaaggtaaccgtag-3 | **Rvs:** 5-aaggtttttattactttgtactctagggc-3 |
| cdtABC_F2 | **Fwd:** 5-tacaaagtaatcaggattagaacatttatcc-3 | **Rvs:** 5-agcttgcatgcctgcaggtctgcaaggggctattccaaagc-3 |
| CiaB1_F1 | **Fwd:** 5-cccggggatcctctagagtctaaataaatgctgataagcttaaag-3 | Rvs: 5-ccttgttacgaaactcatataacgcattatttc-3 |
| CiaB_Kanr | **Fwd:** 5-Atatgagtttcgtaacaaggtaaccgtag-3 | **Rvs:** 5-tttcacaaaattactttgtactctagggc-3 |
| CiaB2_F2 | **Fwd:** 5-tacaaagtaattttgtgaaattgaagataatatttttc-3 | **Rvs:** 5-agcttgcatgcctgcaggtctatttcctataagctcacttac-3 |
| lctP_F1 | **Fwd:** 5-cccggggatcctctagagtcgttatgtgcaatttacaaat-3 | **Rvs:** 5-taaaagtgcagcggttatag-3 |
| lctP_Kanr | **Fwd:** 5-ctataaccgctgcacttttataaccgtag-3 | **Rvs:** 5-aaagcatcacctgtgtgtgcttactttgtactctagggc-3 |
| lctP_F2 | **Fwd:** 5-gcacacacag gtgatgcttt-3 | **Rvs:** 5-agcttgcatgcctgcaggtctaggtacacctttatcttca-3 |
| Δ*lctP*+*lctP* (comp) | **Fwd:** 5-acaccaattgaactaatgattataaccc-3 | **Rvs:** 5-acactctagattgttgaaataaaacttaaa-3 |
| Δ*ciaB*+*ciaB* | **Fwd:** 5-acaccaattggttctcccatatctctcatt-3 | **Rvs:** 5-acactctagaagtcataaaagctcctttgt-3 |
| Δ*cjeN*+*cjeN* | **Fwd:** 5-acaccaattggtttaaaattttctataacat-3 | **Rvs:** 5-acactctagaaatagcaccattaaagtata-3 |
| **Recombinant protein Primers** | | |
| *CjeNc_his6_ | **Fwd:** 5-ctttaagaaggagatatacatatgCAAAATTCTAGTTTTGAAGGAAAAG-3  **Rvs:** 5-agtggtggtggtggtggtgctcgagGAATTTATTGTGTTTTCTCCATTTATAAG-3 | |
| *CjeN_Thr69Ser77Thr111his6_ | **Fwd1:** 5-ctttaagaaggagatatacatatgCAAAATTCTAGTTTTGAAGGAAAAG-3  **Rv**s **1:** 5- tggtgcatctATACCGAAAAAGCTTATTCTAG -3  **Fwd 2:** 5-ttttcggtatAGATGCACCACAACTTAAAC-3  **Rv**s**2:** 5-tttacaatagCAACAATGCTACCATAAGTATC-3  **Fwd 3:** 5-agcattgttgCTATTGTAAAGCTTAATGATGTTG-3  **Rv**s**3:** 5-agtggtggtggtggtggtgctcgagGAATTTATTGTGTTTTCTCCATTTATAAG-3 | |

**Fwd: forward; Rvs: reverse; F1: fragment 1; F2: fragment 2; Comp: Complementation; *Lower case indicate overlap.**

**Table S4: RT-qpCR primers**

| **Primer Name** | **Sequence** |
| --- | --- |
| XP_004336746.1 | Fwd: 5-GCTTGCGTTGGGTCAGT-3  Rvs: 5-CAGCTTGATGTCGTCCTTGT-3 |
| XP_004349665.1 | Fwd: 5-CTACTCCTCAATGTGCTCCAAC-3  Rvs: 5-GCAGTCGGCCATAGTCATTT-3 |
| XP_004334023.1 | Fwd: 5-GAGGAGGAGTTTATGACG-3  Rvs: 5-GAGGGCAATTCCAATCAG-3 |
| XP_004336998.1 | Fwd: 5-AACCAAAGCGAGTCGTACC-3  Rvs: 5-ATCTCCACTCCCTCCTCTTC-3 |
| XP_004352909.1 | Fwd: 5-TACCTCCTCTCCTCGTTCAA-3  Rvs: 5-GAATCGTGTTCGGTTCTCAAAG-3 |
| XP_004346566.1 | Fwd: 5-ACAAGCCCAAGACTCTGATG-3  Rvs:5-TCTGCACCTGGTAGATGGTA-3 |
| XP_004346021.1 | Fwd: 5-CTTTAAGCAGCTCCTCCAGTC-3  Rvs: 5-CCCTCGGTCATGAAGATGTTT-3 |
| XP_004337009.1 | Fwd: 5-AGCGCAATGACAGGTGAA-3  Rvs: 5-AATGCACACCCGGCTAAA-3 |
| XP_004342027.1 | Fwd: 5-TCCGCGAGATCAGGAAGTA-3  Rvs: 5-CTGGAACCTCAGATCAGTCTTG-3 |
| XP_004346618.1 | Fwd: 5- CCACAACATCATTCGCAACC-3  Rvs: 5- GTGCTTGCCGGACTTACA-3 |
| *cdtB* | Fwd: 5-CGCGTTGATGTAGGAGCTAAT-3  Rvs: 5-GTCTTGAAACTGTAGTAGGTGGAG-3 |
| *gyrA* | Fwd: 5-AGTAATACGTGGCACATCAAATTTACTTCTAAT-3  Rvs: 5- GCAGAATTAATGAAAGAAATTGCAAGACTTG-3 |

**Fwd: forward**

**Rvs: reverse**

**Reference**

1.	Nasher, F. *et al.* (2022) Survival of Campylobacter jejuni 11168H in Acanthamoebae castellanii Provides Mechanistic Insight into Host Pathogen Interactions. *Microorganisms* 10. 10.3390/microorganisms10101894

2.	Wang, Y. and Taylor, D. (1990) Natural transformation in Campylobacter species. *Journal of bacteriology* 172, 949-955
